# Pipolins are bimodular platforms that maintain a reservoir of defense systems exchangeable with various bacterial genetic mobile elements

**DOI:** 10.1101/2024.05.22.595293

**Authors:** Víctor Mateo-Cáceres, Modesto Redrejo-Rodríguez

## Abstract

Defense genes gather in diverse types of genomic islands in bacteria and provide immunity against viruses and other genetic mobile elements. Here, we disclose pipolins, previously found in diverse bacterial phyla and encoding a primer-independent PolB, as a new category of widespread defense islands. The analysis of the occurrence and structure of pipolins revealed that they are commonly integrative elements flanked by direct repeats in Gammaproteobacteria genomes, mainly *Escherichia*, *Vibrio* or *Aeromonas*, often taking up known mobile elements integration hotspots. Remarkably, integrase dynamics correlates with alternative integration spots and enables diverse lifestyles, from integrative to mobilizable and plasmid pipolins, such as in members of the genera *Limosilactobacillus*, *Pseudosulfitobacter* or *Staphylococcus*.

Pipolins harbor a minimal core and a large cargo module enriched for defense factors. In addition, analysis of the weighted gene repertoire relatedness revealed that many of these defense factors are actively exchanged with other mobile elements. These findings indicate pipolins and, potentially other defense islands, act as orthogonal reservoirs of defense genes, potentially transferable to immune autonomous MGEs, suggesting complementary exchange mechanisms for defense genes in bacterial populations.

## INTRODUCTION

Prokaryotic genomes are constantly influenced by foreign mobile genetic elements (MGEs), which directly affect genomic plasticity, adaptation, and evolution^1,2^. MGEs range from a single transposase gene that facilitates its own transposition within the same molecule^3^ (*i.e.* Insertion Sequences or IS) to large genomic sequences that constitute complex biological systems such as bacteriophages^4^ and conjugative elements^5,6^. Moreover, the mobilization machinery of an MGE can induce modifications in the host genome, granting the MGE an adaptive role of its own^7^. However, the primary source of selective advantages in complex MGEs is the presence of cargo genes or genes not directly involved in their replication, mobilization, or transference. These genes are often associated with antimicrobial resistance^8^ (AMR) and virulence factors^9^, providing significant benefits to the host organism. Additionally, recent studies have shown that MGEs can act as both targets and vectors for bacterial defense systems^2,10^, simultaneously hijacking and restricting horizontal gene transfer^11,12^.

Acquisition of advantageous traits through MGE transference is the main form of short-term adaptation in bacteria, critical for cell survival in the context of infection and antimicrobial treatment^13^. The direct implication of MGEs in bacterial survival has spurred intensive research aimed at understanding the emergence of multi-drug resistant (MDR) bacteria and the determinants of phage resistance, essential for advancing modern phage therapy^14^. This research has led to the expansion MGE classes, including Integrative and Conjugative Elements (ICEs)^6,15^, integrons^16^, CRISPR-Cas associated transposons^17^ (CASTSs) or tycheposons^18^. Among the new types of MGEs discovered, *pipolins* stand out for their extensive genetic diversity and variability^19,20^. Pipolins comprise a group of MGEs distinguished by encoding a replicative DNA polymerase from the B family (PolB) and can be therefore referred to as “self-replicating” elements^21^, together with other PolB-enconding MGEs like polintons^22^ and casposons^23^. Unlike the PolBs of polintons and casposons, PolBs encoded in pipolins have been show not to require a preexisting primer to initiate the synthesis of the complementary strand, hence they are referred to as *primer-independent* PolBs or piPolBs^19^.

Interestingly, piPolBs have been identified in diverse bacteria and several mitochondria, suggesting an ancient origin predating the divergence of major bacterial phyla. Phylogenetic analysis revealed that piPolBs can be grouped into two main clades congruent with their host: Gram-positive and Gram-negative piPolB, with the latter group also including mitochondrial piPolBs. However, polymerase phylogeny was inconsistent at lower levels with the host evolution, suggesting an extensive horizontal transfer of the piPolB characteristic of genes from MGEs^19^. Accordingly, a survey of pathogenic *Escherichia coli* strains showed that pipolins can be transferred between strains across diverse phylogenetic groups and pathotypes^20^.

Alongside the piPolB gene, previously known pipolins typically encode a recombinase gene, which might enable the integration and excision of the pipolin into a tRNA gene by means of flanking att-like direct repeats (DR)^19^, as well as a variety of genes related to DNA metabolism such as restriction-modification (RM) systems, helicases, exonucleases, ribonucleases, AAA ATPases and HTH/Zn-finger DNA binding proteins. Although pipolins are mainly integrative elements, piPolB genes have been detected in circular plasmids of *Staphylococcus epidermidis* (pSE-12228-03), *Lactobacillus fermentum* (pLME300) and the mitochondria of the fungus *Cryphonectria parasitica* (pCRY1)^19^, increasing the genetic structure, diversity and lifestyles of pipolins. Like other MGEs, unknown function genes are also frequent, hindering our understanding of the biological significance of pipolins. Unexpectedly for such a widespread element, no AMR gene or virulence factor has yet been linked to pipolins, leaving the piPolB as the sole pipolin hallmark. However, the persistence of pipolins in distant taxa, along with recent evidence of horizontal gene transfer events in *E. coli*, suggests that these are ancient elements providing biological traits that offset the fitness cost associated with their maintenance^24,25^.

In this study, we conducted an extensive screening of bacterial genomes from the NCBI Assembly database to assess the prevalence of Pipolins. Our analysis covered over one million bacterial genome assemblies, significantly expanding the scope and diversity compared to previous studies. As a result, we identified novel divergent piPolB groups and pipolins in species not previously reported, including significant pathogens such as *Salmonella enterica* and *Staphylococcus aureus*. We identified and analyzed pipolins and piPolB-containing sequences in more than 11,000 different assemblies, revealing diverse integrase shuffling patterns and novel integration sites associated to pipolins. Comparative analysis of genes encoded in pipolins along with a comprehensive dataset of plasmids, conjugative and integrative elements (ciMGEs), and bacterial viruses revealed that these elements serve as active platforms for the transfer of various genetic systems, with defense factors being by far the most abundant. To further understand the interactions between pipolins and other MGEs, we analyzed the weighted gene repertoire relatedness (wGRR) shared by these elements. Our analysis revealed that pipolins show extensive recombination with ciMGEs and plasmids, particularly highlighting the exchange of defense genes in enterobacteria. Altogether, we propose that pipolins may be the paradigm of a group of MGEs specialized in defense functions that would offer a variety of immune genes to be incorporated by autonomous genetic mobile elements.

## MATERIALS AND METHODS

### Pipolin Screening in Genbank Assemblies

All bacterial genome assemblies from the GenBank database (1,310,495 genomes, accessed on November 11, 2022) were downloaded using the NCBI Datasets command line tool (version 13.43.2, https://www.ncbi.nlm.nih.gov/datasets) with the “--exclude-atypical” option and analyzed for the presence of pipolins using ExplorePipolin^26^. Briefly, ExplorePipolin is a pipeline that analyzes the presence of pipolins in multi-fasta nucleotide sequence files and returns several files describing pipolin genome coordinates, flanking direct repeats, genes encoded in the element, and other useful information. Pipolins are detected using a piPolB HMM profile and a reference direct repeat (DR) sequence from *E. coli* 3-373_03_S1_C2 to delimit pipolin boundaries. Additionally, if DRs are not detected by sequence similarity, ExplorePipolin conducts a *de novo* DR search where repeats are searched by aligning with blastn both sides of the piPolB genetic context. *De novo* DRs are validated if they overlap a tRNA or tmRNA gene. If this method fails to find valid DRs, ExplorePipolin cuts 30 kbp at each side of the piPolB gene and the pipolin is denoted as “minimal” or “incomplete”. Genomes lacking piPolB genes were discarded from further analysis.

After ExplorePipolin execution, complete pipolins exceeding 100 kbp were also cut at 30 kbp from each piPolB flank since we do not expect pipolins to exceed that length according to previous studies^19,20^. Also, if a pipolin showed inconsistencies between reconstruction versions (*i.e.* different pipolin lengths between versions), the reconstruction was omitted and a 30 kbp flanking window was extracted from the contig encoding the piPolB.

### piPolB Phylogenetic Analysis

A total of 9,740 piPolB amino acid sequences longer than 800 amino acids were obtained from the ExplorePipolin result files. Sequences shorter than 800 amino acids were not included in the analysis since partial sequences could hinder the phylogenetic inference. Bam35 DNAP (Uniprot: Q6X3W4) was selected as an outgroup. Multiple sequence alignments were calculated using MAFFT-L-INS-i^27^ and trimmed with trimAI^28^, both with default parameters. Phylogenetic inference of the polymerases was carried out by IQ-TREE^29^, which relies on ModelFinder^30^ to choose the best evolutive model. The best-fit model was Q.pfam+I+I+R10 according to the BIC (Bayes Information Criteria) value. The number of ultrafast bootstrap replicates and the number of SH-like approximate likelihood ratio test (SH-aLRT) replicates were set at 1,000.

### Pipolin reannotation and candidate integrase search

To improve pipolin annotation we re-predicted, clustered, and annotated the coding sequences with several benchmarked tools. Firstly, proteins encoded in pipolins were predicted by Prodigal^31^ (V2.6.3) using “-c” in order to include plasmid genes that sometimes are interrupted by the sequence ends in FASTA files. The resultant 316,154 protein sequences were co-clustered with representative sequences from a database of annotated MGE proteins (mobileOG-db^32^, 68,919 sequences after running CD-HIT^33^ with 70% identity limit). Clustering was carried out using MMseqs2^34^ (Version: 15.6f452) with the following parameters: –min-seq-id 0.35 -c 0.6 –cluster-mode 0 -s 7.5 –cluster-steps 9. Sequences shorter than 30 amino acids and longer than 2000 amino acids were removed from the clustering process. Then pipolin protein function was inferred both at individual level, using EggNOG-mapper v2^35^ for a general annotation, and at cluster level by taking representatives with CD-HIT (-c 0.9), aligning them with MAFFT and running hhblits 3.3.0^36^ against the Pfam35^37^ database (-n 5 iterations, top 3 hits with E-value < 0.001 kept). Additionally, we used specific profiles to annotate relaxases (through MOBScan^38^), integrases^39,40^, and piPolBs^26^, applying the methods and thresholds used in the reference articles.

After clustering and annotating pipolin genes, we created a binary presence-absence matrix where each row represents a piPolB gene and each column represents a gene cluster. In this matrix, entries are assigned a 1 if there is a gene from a cluster encoded within the piPolB genetic context (*i.e.* the pipolin or the 30 kbp window) or a 0 if there are no genes from a cluster near the piPolB. To find candidate integrases associated with piPolBs, for each cluster of piPolBs with size of at least 40 sequences (which correspond to the main piPolB clades in the phylogeny) this matrix was split into two submatrix, M_A_ and M_B_, containing M_A_ all rows where the piPolB cluster is present and M_B_ all rows where the piPolB are not present. Next, in each submatrix, we calculated the frequency of observing a recombinase cluster *R* whenever the piPolB cluster is present (f_RcMA_) or no (f_RcMB_). Then, we considered as a candidate piPolB-associated recombinase, all recombinases that showed a differential observed frequency (Δf_R_ = f_RcMA_ - f_RcMB_) higher than 0.05 in one or more piPolB clusters. Finally, all candidate integrases found in pipolins where no DR could be detected, were manually checked for the presence of adjacent tRNAs or interrupted ORFs within a conserved genetic context that allowed a confident delimitation of the pipolin.

### MGE comparative characterization

To carry out a comparative characterization of pipolins we assembled a dataset composed of plasmids, integrative conjugative and mobilizable elements (ciMGEs), phages and pipolins. After the removal of duplicated sequences and elements shorter than 5 kbp or longer than 500 kbp, the final dataset comprised 50,022 plasmids from PLSDB^41^ (version: 2023_11_03_v3), 10,925 ciMGEs from a recent study^42^ complemented with CIMEs (cis-integrative mobilizable elements) from the ICEBerg 3.0^43^ database, 10,925 phages from the PhageScope^44^ database (only phages derived from RefSeq, GenBank, EMBL, and DDBJ were selected), and 7,409 pipolins from this study, after removal of duplicated sequences and pipolins shorter than 5 kbp or longer than 500 kbp. Pipolins were also re-delimited if a new alternative integration site was described (see Results) and pipolins lacking any defined boundaries were further trimmed to 10 kbp to each side of the piPolB, concordant with the mean size of these MGEs. Pipolins lacking a piPolB of at least 800 aminoacids or encoding less than 8 genes were also not included.

To ensure a homogeneous annotation that allowed comparisons between MGE families, we predicted plasmid, ciMGE and phage genes using Progidal (V2.6.3) with the same parameters used previously with pipolins. Then, annotation of phage-like functions was carried out using PHROGs^45^ profiles and conjugation functions were annotated using CONJScan^46^ profiles. These profiles were used by HMMER 3.4^47^ (hmmsearch), and we kept the best results with at least 0.001 E-value and 50% coverage. Genes conferring adaptive traits were annotated with specialized tools: AMR genes were detected by AMRFinderPlus 3.12.8^48^, defense systems were identified by PADLOC v2.0.0^49^ and virulence factors were determined by aligning MGE CDSs to the VFDB^50^ protein sequences (full dataset, accessed on 07.03.2024) using MMSeqs2 and keeping the best result showing at least 80% sequence identity and 80% coverage.

### Identification of recent gene exchange events between MGEs

The original MGE dataset was filtered to keep only MGEs that may have exchanged genetic material with pipolins. To this end, all protein sequences were clustered at 75% identity and 75% coverage using linclust from MMSeqs2. Then, MGEs are kept if they have at least one gene in a cluster containing sequences from at least three pipolins and make up at least 3% of the MGEs represented in the cluster. Largest clusters where more than 2000 sequences are represented were discarded as they only contain ubiquitous IS transposases or short HTH-containing proteins that would bias the study.

Next, we adapted a recently used method^51^ for the detection of “recombining genes” (RGs) between plasmids, phages and plasmid-phages^52^ (P-Ps). Briefly, this method consists of identifying highly similar pairs of proteins encoded in very different MGEs as measured by their weighted gene repertoire relatedness (wGRR). The wGRR is defined by the sum of protein sequence identities for each best-bidirectional hit (BBH) between two elements, A and B, divided by the number of genes of the smaller element:

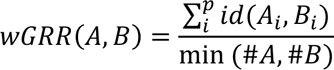

Thus, the wGRR between two MGEs is directly proportional to the similarity of the elements and ranges between 1 (identical elements) and 0 (unrelated elements). Due to the computational cost of computing an all-vs-all protein comparison, we reduced the number of alignments by performing an initial clustering at 35% sequence identity and 50% coverage with cluster from MMSeqs2. All-vs-all protein alignment were then computed inside each cluster with MMSeqs2 prefilter (-c 0.5 -s 7.5 –max-seqs N=cluster size) and align (-a -c 0.5 -e 0.0001 --min-seq-id 0.35).

RG determination was performed by identifying BBH with elevated sequence similarity (>80% identity and >80% coverage) found in MGEs sharing a low wGRR value (< 0.2). If the number of RGs between two MGEs exceeded 25, these genes were not labeled as RGs as they may be the observation of a cointegrated MGE rather than actual exchanges. In the original work, authors used a wGRR threshold of 0.1 for RG determination with satisfactory results, but due to the reduced conserved core of pipolins we needed to raise the wGRR limit to 0.2 (MGE-pipolin exchanges may comprise more than 10% of the pipolin, see Results). Furthermore, the 0.2 threshold is within the 0-0.3 wGRR range where highly similar BBH explained by recent exchanges^53^. Genes not labeled as RGs were classified as non-recombining genes (NRGs), while genes lacking a homolog (*i.e.,* genes that are not present in the BBH list) are classified as NRG with no homologs (nhNRGs). Note that genes are classified as RG, NRGs or nhNRGs in the context of our dataset, meaning that the identification of a gene as NRG does not confirm that it has never been exchanged. Studies with alternative datasets may entail the detection of more RGs not found in this work.

Quantification of genetic exchanges between MGEs and the statistical comparison between RGs and NRGs was performed following the methods described in the original article. Briefly, quantification of exchanges required an initial grouping of highly similar RGs (> 80% identity and >80% coverage) in families using a single-linkage algorithm. Then, a single exchange event is counted for each cluster, which is classified according to the MGE classes observed in the cluster. The statistical analysis required the assignment of functional categories to RGs and NRGs, for which PHROG^45^ and VDFB^50^ classes, CONJScan^46^ genes, PADLOC^49^ defense systems and top PFAM hits (derived from the clustering and reannotation of pipolins) were used. Then, over- and underrepresentation of genes in each functional category was tested in contingency tables using the exact Fisher’s test with the Benjamini-Hochberg multiple testing correction^51^.

## RESULTS AND DISCUSSION

### 1. Patchy pipolin prevalence across all major bacteria groups

In order to understand the diversity and evolution of pipolins, we performed a complete screening of the bacteria assemblies in GenBank Datasets, using the tool ExplorePipolin, based in profile searches against a piPolB curated HMM profile^26^. We detected putative pipolins in 11,431 of >1.3M genome assemblies, comprising a prevalence of nearly 0.9% of bacterial genomes (**Figure 1**). Assemblies from *Gammaproteobacteria*, where *Escherichia* (67.2%), *Vibrio* (14.5%), *Aeromonas* (1.1 %) and various enterobacteria genus like *Enterobacter* (3.1%), *Salmonella* (1 %) or *Citrobacter* (1%), make up most of the largest part of our pipolin dataset. However, other genera from diverse distant taxa are well represented: *Pseudosulfitobacter* (1.2 %), *Staphylococcus* (1.1%), *Limosilactobacillus* (0.8%), *Corynebacterium* (0.7%), highlighting the presence of pipolins in all major bacterial clades. Although this database is biased towards medically relevant species, our comprehensive screening unveiled a notable prevalence of pipolins in some relevant genera. For instance, in *Vibrio*, pipolins were found in 9.7% of the genome assemblies, while in *Aeromonas* and *Limosilactobacillus*, the prevalence was even higher at 12% and 14%, respectively.

**Figure 1.**
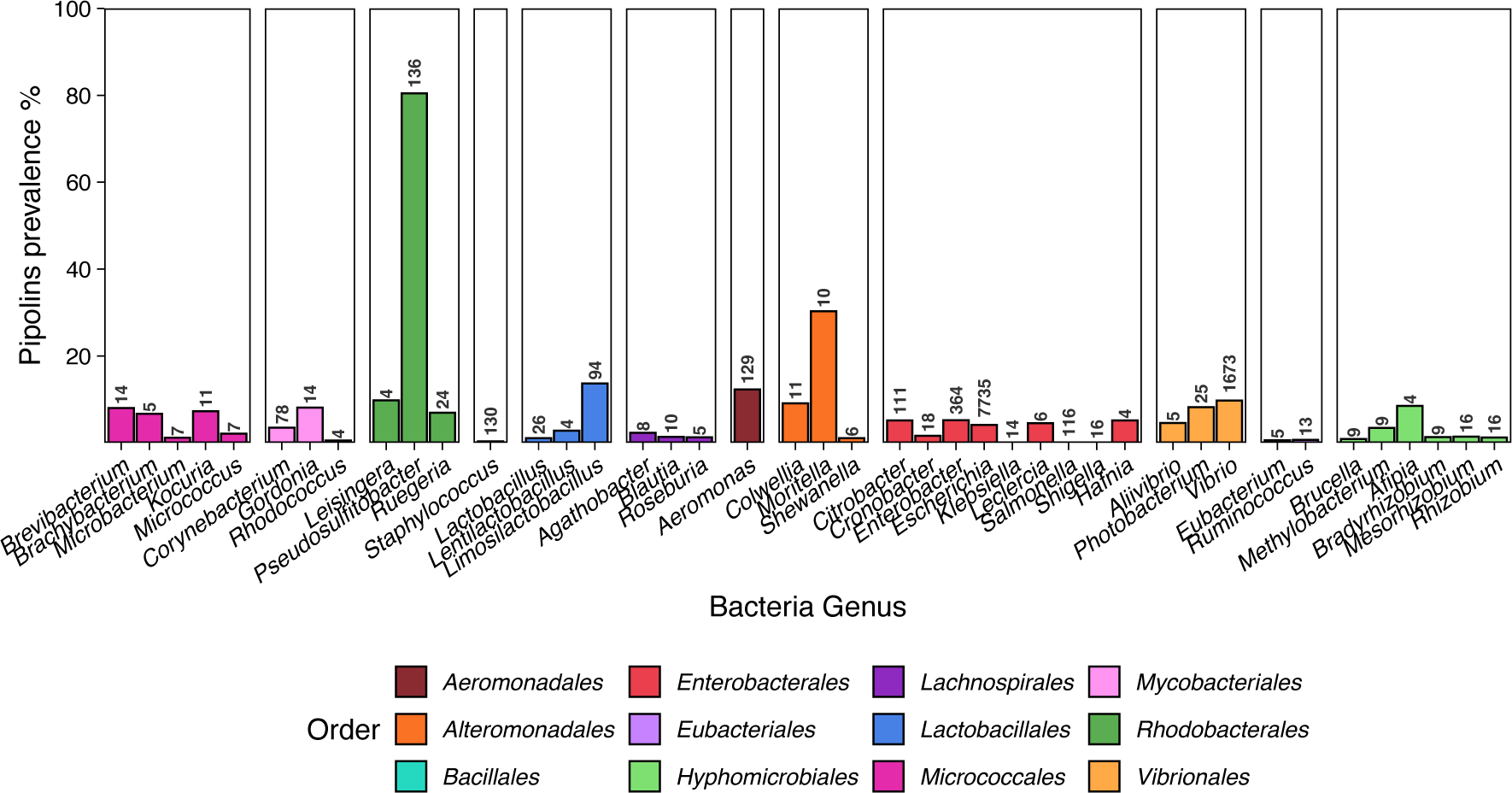
Distribution of pipolin in bacterial genome assemblies. The prevalence of pipolins is represented by the ratio (%) of pipolins to the total genome assemblies per genus in the Genbank database grouped by order. The total number of pipolins in each genus is also shown. Only genera with more than 3 detected piPolB-containing sequences are shown.

In total, we detected 11,714 pipolins from 11,431 genome assemblies, congruent with the presence of several piPolBs in some genome assemblies (**Figure 2A**). Pipolins were present in assemblies from 181 distinct genera and 397 distinct species (**Supplementary Dataset 1**). Besides previously known pipolins from opportunistic pathogens like *E. coli* or *S. epidermidis*, we confirm the presence of this MGE in widely known pathogens from diverse phyla, some of them not previously reported, such as *Vibrio cholerae*, *Salmonella enterica*, *Staphylococcus aureus* and *Corynebacterium diphtheriae*. Phylogenetic analysis of the piPolBs revealed two main clades related to the host phylogeny. The Gram-negative piPolBs can be subdivided in two subgroups: a main group comprising *Gammaproteobacteria* (includes *Vibrio*, *Aeromonas*, *Escherichia*, *Enterobacter*, and other enterobacteria) and a minor group including *Alphaproteobacteia* (*Rhizobiales*, *Rhodobacteriales*, etc.). Similarly, Gram-positive piPolBs involve several subgroups: *Clostridia*-*Actinobacteria*, *Lactobacillaceae*, and *Staphylococcus*. This result is congruent with the first search for piPolBs carried out in previous studies^19^. However, at lower taxonomy levels, we observed multiple inconsistencies with the host evolution (**Figure S1**) that suggested more recent events of horizontal gene transfer. Moreover, the presence of nearly identical piPolBs in diverse enterobacteria indicates these events may transcend the genus barrier. We also observed strong discrepancies between piPolB and host phylogeny inside the *Lactobacillaceae* family and between different staphylococcal species. Surprisingly, some assemblies from *Aeromonas*, *Limosilactobacillus*, and especially, *Vibrio* encode more than one distinct piPolB, with sequence identities around 35%. This suggests that pipolin evolution in these genera has reached a point where the element could have duplicated and diverged through evolutionary processes.

**Figure 2.**
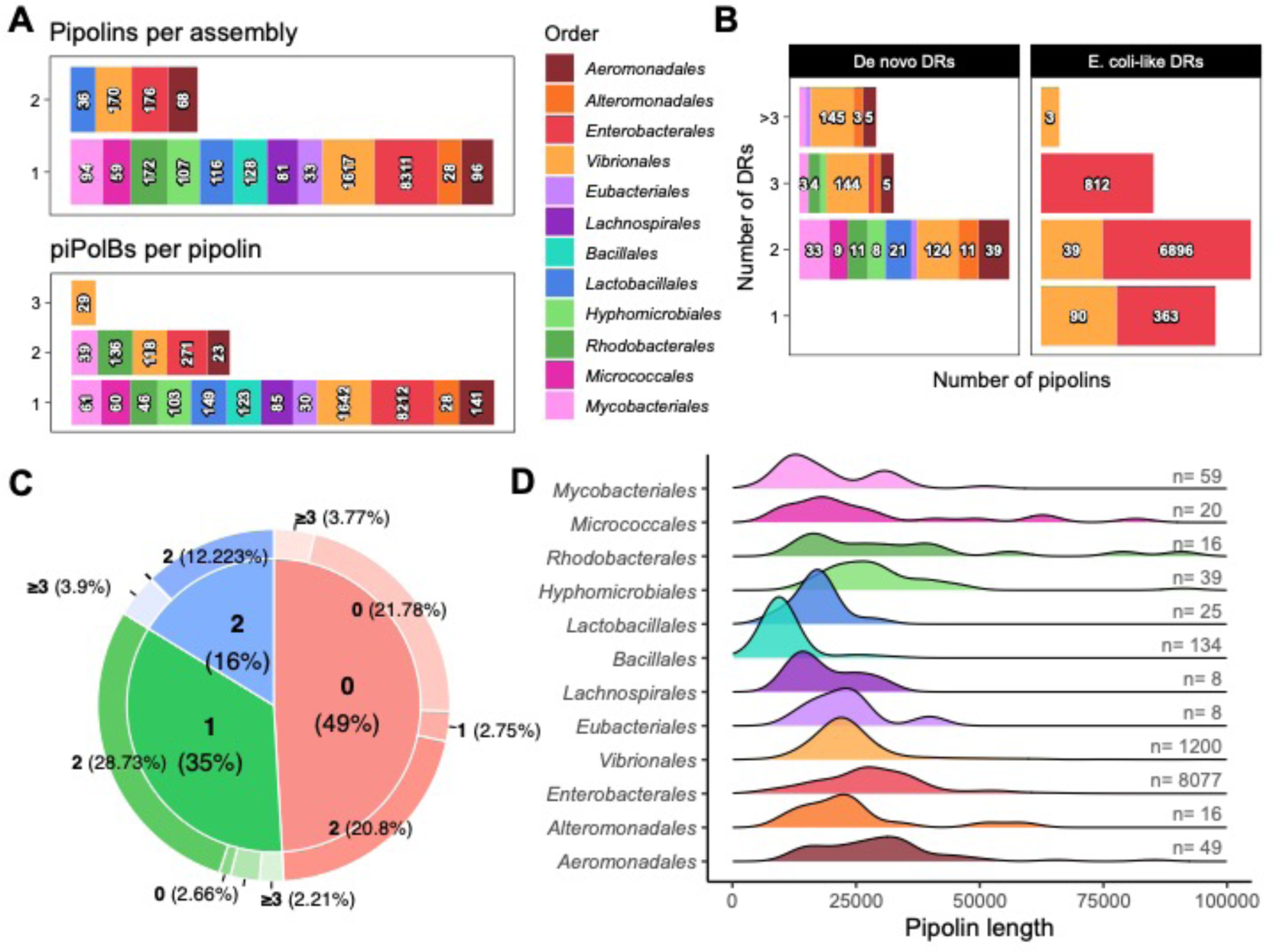
Pipolins reconstruction and structure. **A.** Number of pipolins detected per assembly and piPolB genes per pipolin. Only orders with more than 20 putative pipolins were included. **B.** Presence of att-like direct repeats in pipolins (DR). Number of *de novo* detected (left) or known (right) direct repeats in each pipolin. Numbers are shown above stacked bars scaled logarithmically and colored by bacterial order. **C.** Ratio of pipolin reconstruction gaps (inner piechart) and number of detected direct repeats (DRs, outer circle). **D.** Length of pipolins by bacterial order. Only reconstructed integrative pipolins with DRs or alternative integration sites (see text), and plasmid-pipolins were considered. The number of delimited pipolins in each order is indicated.

### 2. Pipolins flexibility as revealed by diverse associated integrases

#### 2.1. Diversity of pipolins structure and boundaries

Using ExplorePipolin^26^ we were able to extract 8,646 *bona fide* complete integrated pipolins in bacteria assemblies, where the known direct repeats (DR) and adjacent tRNA integration site were successfully predicted, being around 6,000 the most-likely reconstruction from different contigs (**Figure 2B-C**). Additionally, it annotated 2,896 varied incomplete pipolins or piPolB-containing fragments where known flanking DRs and/or integration sites could not be identified due to deletions near the ends of the element, the presence of alternative structures (e.g., circular plasmids), or integration into a site other than a tRNA, as discussed below.

As expected, most of complete pipolins were found in *E. coli* and other *Enterobacteria.* In contrast, *de novo* DR detection enabled the prediction of complete pipolin with delimited boundaries in other distant taxa such as *Corynebacterium* and *Limosilactobacillus* (**Figure S1, Supplementary Data 1**). Analysis of complete pipolins revealed that they can integrate into a variety of tRNA genes and even tmRNAs, which is largely related to the piPolBs clade and, overall, is contingent on the recombinase encoded in the element (see below). Besides, pipolin length shows a distribution centered around 20 kbp, suggesting these are short MGEs compared to phages, conjugative plasmids and ICEs and more similar to mobilizable and parasitic elements (**Figure 2D**). However, mean pipolin length is highly dependent on host as can be seen in *Vibrio* or *Aeromonas*, where Pipolins are shorter compared to *Escherichia* and other *Enterobacteria*.

Reconstruction and delimitation of incomplete pipolins (piPolB-containing fragments), besides possible assembly artifacts, the nearby presence of adjacent prophages or Insertion Sequences (IS) increases difficulty in both sequencing and pipolin extraction. Furthermore, incomplete pipolins could correspond also to plasmid pipolins lacking an integration mechanism as in the case of pLME300 from *Lactobacillus fermentum*^54^, pSE-12228-03 from *Staphylococcus epidermidis* and other staphylococcal plasmids found in this screening (see below).

Although complete Pipolins make up around 75% of our dataset, many of them belong to species phylogenetically close to *E. coli* while pipolins from distant species often lack any DR (**Figure 2A-C**). Therefore, we analyzed integrases co-located with piPolBs with the aim of finding new alternative integration sites for non-canonic pipolins. This allowed us to delimit more pipolins and carry out a wider analysis of pipolin genetic content. Specifically, we clustered all the predicted CDSs at 35% identity sequence and 60% coverage using mmseqs2^55^ (**Supplementary Dataset 2**) and built a presence-absence matrix where rows represent piPolB genes ordered by their phylogeny and columns represent integrase clusters. Then, we selected as candidate integrases those clusters showing a differential observed frequency of 0.05 or greater. This is the frequency of observing an integrase cluster and a piPolB cluster in the same pipolin minus the frequency of observing that integrase cluster in pipolins with other piPolB clusters (see Methods). In total, 28 integrases clusters, 15 tyrosine recombinases (YR) and 13 serine recombinases (SR), were present next to at least 7 different piPolB clusters and were manually revised to find an integration site associated with the recombinase (**Supplementary Dataset 3**).

#### 2.2. Most abundant pipolins in *Gammaproteobacteria* derive from a common ancestor

We found 6 recombinase clusters near *Gammaproteobacteria* piPolBs related to previously described families of ICEs, genomic islands, or phage recombinases^40^ (**Figure 3A**). These recombinases belong to YRs from the Int_SXT_ (two clusters) and Int_P2_ (four clusters) families. Int_SXT_ recombinases are ubiquitously found in *Gammaproteobacteria* pipolins and usually encoded near the integration site (**Figure 3B**). Synteny analysis between distant elements revealed that, besides the piPolB gene, the Int_SXT_ is usually surrounded by a gene encoding for a DUF2787, a WYL domain-containing protein and several protein orphans. Notably, DUF2787 can be found in the *Vibrio cholerae* seventh pandemic island II (VSP-II)^56^, and WYL domain is a nucleic acid sensor involved in defense and DNA damage responses^57,58^. This synteny preservation observed in distant pipolins and the low sequence identity shared by these genes (around 30-35% for piPolBs and YRs from *Escherichia* and *Vibrio*) indicates a robust evolutive association. Furthermore, element comparison also revealed that an Int_P2_, usually followed by an excisionase and located near the piPolB, is present in all *Enterobacterales* and closer *Aeromonadales* and *Vibrionales* (**Figure 3B**, *pipolins 1 to 5*). However, unlike the Int_SXT_ that is present in all the complete pipolins in *Gammaproteobacteria,* the Int_P2_ recombinase is missing in distant *Vibrionales* and certain *Aeromonadales* subgroups (**Figure 3B**, *pipolins 6 and 8*) suggesting it may have played a secondary role in pipolin diversification while the Int_SXT_ would be involved in the integration and excision of the element. Lastly, a less frequent group of Int_P2_ YRs are encoded in prophages integrated adjacent to the pipolin and probably share mechanism and integration site^20^. Interestingly, a considerable number of genome assemblies from *Enterobacteria* contain pipolins showing a deletion between the Int_SXT_ integrase and the piPolB as well as frequent sequencing gaps compatible with the presence of IS near the Int_P2_, generating the prediction of a truncated version of the YRs (**Figure 3B***, pipolin 2*). Furthermore, piPolBs encoded in this subgroup of pipolins conform a monophyletic clade, suggesting a common origin for this truncated version of the pipolin, and show frequent cointegration of adjacent prophages. Disclosing the relationship between pipolin deletions and prophage cointegration will require further studies.

**Figure 3.**
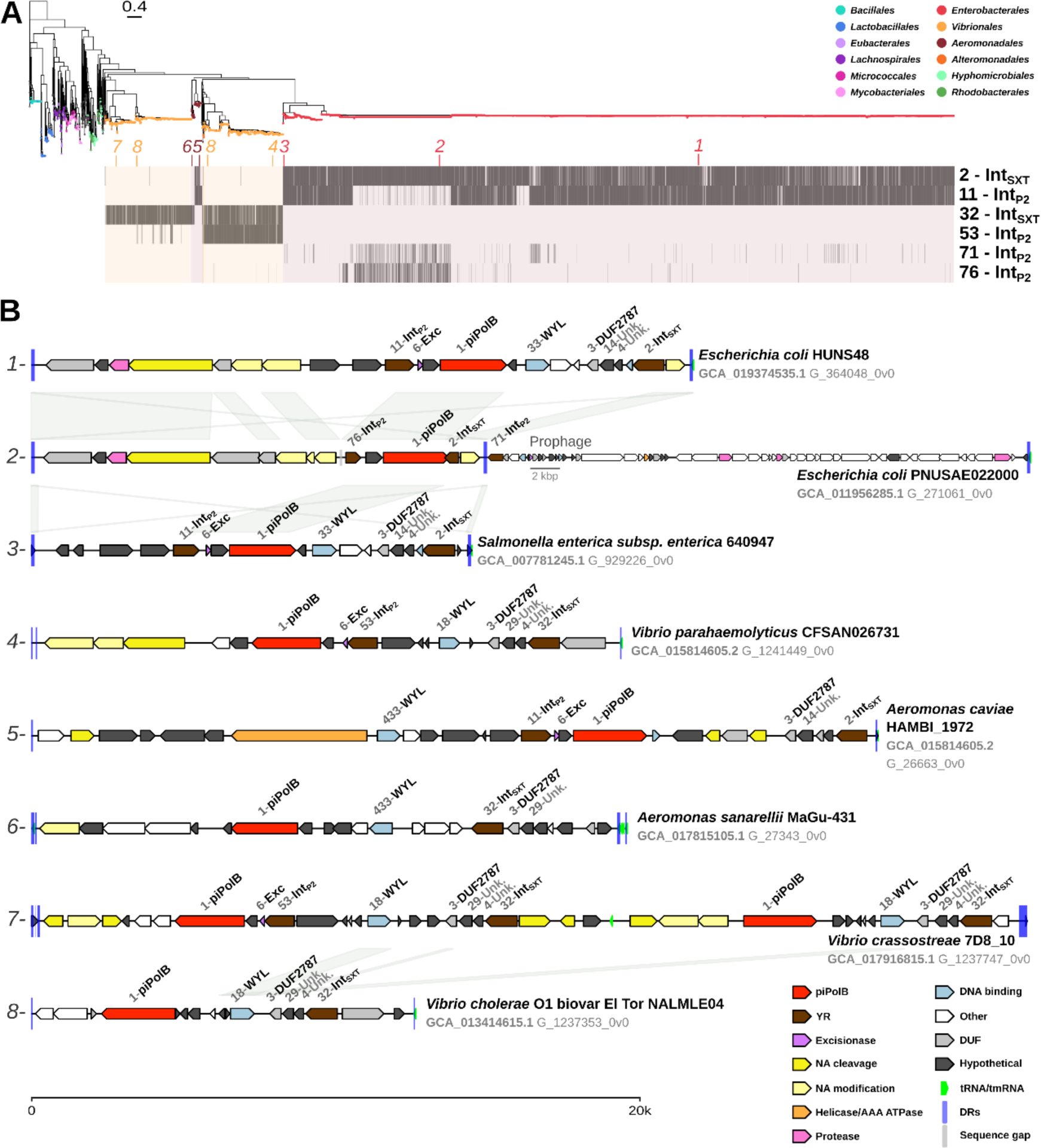
Pipolin integrases in *Gammaproteobacteria*. **A**. Maximum-likelihood phylogenetic tree of 9,740 piPolBs and Bam35 DNAP (outgroup) inferred by IQ-TREE v2 (see Methods). ModelFinder best-fit model was Q.pfam+I+I+R10 according to BIC. Colored tips represent the most frequent orders as indicated. The scale bar indicates substitution rate per site. The heatmap under the tree indicates the presence (black) or absence (empty, clade color) of each integrase cluster within the piPolB genetic context (*i.e.* the pipolin or the pipolin-containing sequence). Numbers next to integrase type corresponds to cluster number (see **Supplementary Data 4**). Italic numbers over the heatmap indicated the location of the example pipolins represented below. **B**. Genomic organization of representative pipolins. Predicted protein-coding, tRNA, and tmRNA genes are represented by arrows, indicating the direction of transcription. Direct repeats (DRs) and sequence gaps are represented by blue and grey bars, respectively. Integrases and other core pipolin genes are labeled indicating cluster number and annotation. Genes are colored according to general functions explained in the legend. Links between genes indicate highly similar regions between pipolins as calculated by minimap2^60^ with -X -N 50 -p 0.1 parameters. Each pipolin is labeled indicating strain name, genome accession number and pipolin identifier, according to ExplorePipolin and screening nomenclature (G_+genome number_+pipolin number_+version). Of note, genes belonging to clusters 14 and 29 are homologues, similarly to Int_SXT_ (clusters 2 and 32), Int_P2_ (clusters 11 and 53), and WYL (clusters 18 and 33).

Apart from cointegrated prophages, the presence of a few pipolins containing Int_P2_ in the group of *Vibrionales* pipolins containing only Int_SXT_ led us to the unexpected identification of systems of two adjacently pipolins integrated into the same tRNA gene (**Figure 3B***, pipolin 7*). In these cases, we found that one element belonged to the group of pipolins containing the two integrases (Int_SXT_ and Int_P2_) while the other element belonged to the Int_SXT_-only group. Both the Int_SXT_ and the piPolB from each individual pipolin show relatively low percentage of amino acid sequence identity (around 30-35%), discarding the possibility of a recent duplication event. Rather than that, the observation of similar individual pipolins in other *Vibrio* species supports the idea that pipolins can be horizontally transferred to other pipolin-containing strains and integrate into the same target gene. These “reinfection” events are probably facilitated by the high divergence of their integrases, allowing them to target different motifs near the tRNA. Furthermore, since we did not detect any pipolin reinfection in *Enterobacterales* we considered that pipolins reached this order relatively later than *Vibrionales* or *Aeromonadales* and their integrases have not diverged enough to colonize alternative integration spots. This idea is supported by the diversity differences observed in pipolins from *Enterobacteriales*, where they show higher synteny and identical integration mechanism, while pipolins from *Vibrionales* or *Aeromonadales* show Int_P2_ deletions or rearrangements in certain subgroups. Tandem accretion leading to element deletions and rearrangements has already been reported for ICEs and other genomic islands^59^, so pipolins would not be an exception. Thus, we propose that pipolins in *Enterobacteriales* were acquired more recently, likely from a *Vibrionales* or *Aeromonadales* common ancestor that contained a pipolin belonging to the Int_SXT_+Int_P2_-Exc group.

Taken together, these results show that piPolBs in *Gammaproteobacteria* are strongly associated to a YRs of the Int_SXT_ family, as well as a DUF2787 and WYL domain-containing genes. Therefore, in agreement with the piPolB phylogeny, it is likely that pipolins in *Gammaproteobacteria* derived from a common ancestor element which diverged in the different variations observed in this study. The IntP2 would play a secondary role and could possibly have been incorporated into the pipolins from a prophage.

#### 2.3. High diversity of pipolins structure and lifestyles beyond *Gammaproteobacteria*

Unlike Gammaproteobacteria, pipolins from other groups exhibit a broader range of genetic structures and compositions (**Figure 4**). In *Alphaproteobacteria*, piPolBs are usually near a large serine recombinase (LSR), which would be responsible for the integration of the pipolin similarly to other MGEs^39^; and a shorter SR (SSR), related to resolvases involved in the maintenance of the circular form of the element (**Figure 4***, pipolin 1*). In the absence of DRs and a tRNA integration sites, we decided to analyze the genetic context of this recombinases, which revealed that these pipolins are integrated into or next to the *yifB* gene, a Mg^2+^ chelatase previously reported as integration site for SRs^61^. Several groups of pipolins in *Alphaproteobacteria* also contain a YR gene from the Int_P2_ or Int_Tn916_ families. Strikingly, while in the first case it has no apparent effect on the integration site, remaining as dependent on the LRS (**Figure 4***, pipolin 2*), the presence of a Int_Tn916_ is at the expense of the LSR, allowing the pipolin to integrate into a tRNA gene and the generation of two DRs similarly to enterobacterial pipolins (**Figure 4***, pipolin 3*). Interestingly, recombinase replace events are not restricted to *Alphaproteobacteria* as pipolins belonging to Gram-positive bacteria also contain diverse YRs and LRSs that are exchanged at different points in pipolin evolution. In *Actinomycetota* (formerly *Actinobacteria*), the presence of a YR from the Int_Tn916_ class or different LSRs is represented by a mutual-exclusion pattern strongly incongruent with the piPolB phylogeny and indicates exchange events as in *Alphaproteobacteria*. Likewise, pipolins encoding Int_Tn916_ are integrated into a tRNA-gene and their DRs could be detected (**Figure 4***, pipolin 4*). Analysis of the genetic context allowed us to establish the *ychF* or the *Ftsk/SpoIII* genes as new integration sites for pipolins encoding distinct groups of LSRs (**Figure 4***, pipolin 5 to 7*). The presence of resolvase-like SRs in pipolins already encoding a large recombinase supports the idea that pipolins can be circularized for replication and employ resolvases to handle replication intermediates.

**Figure 4.**
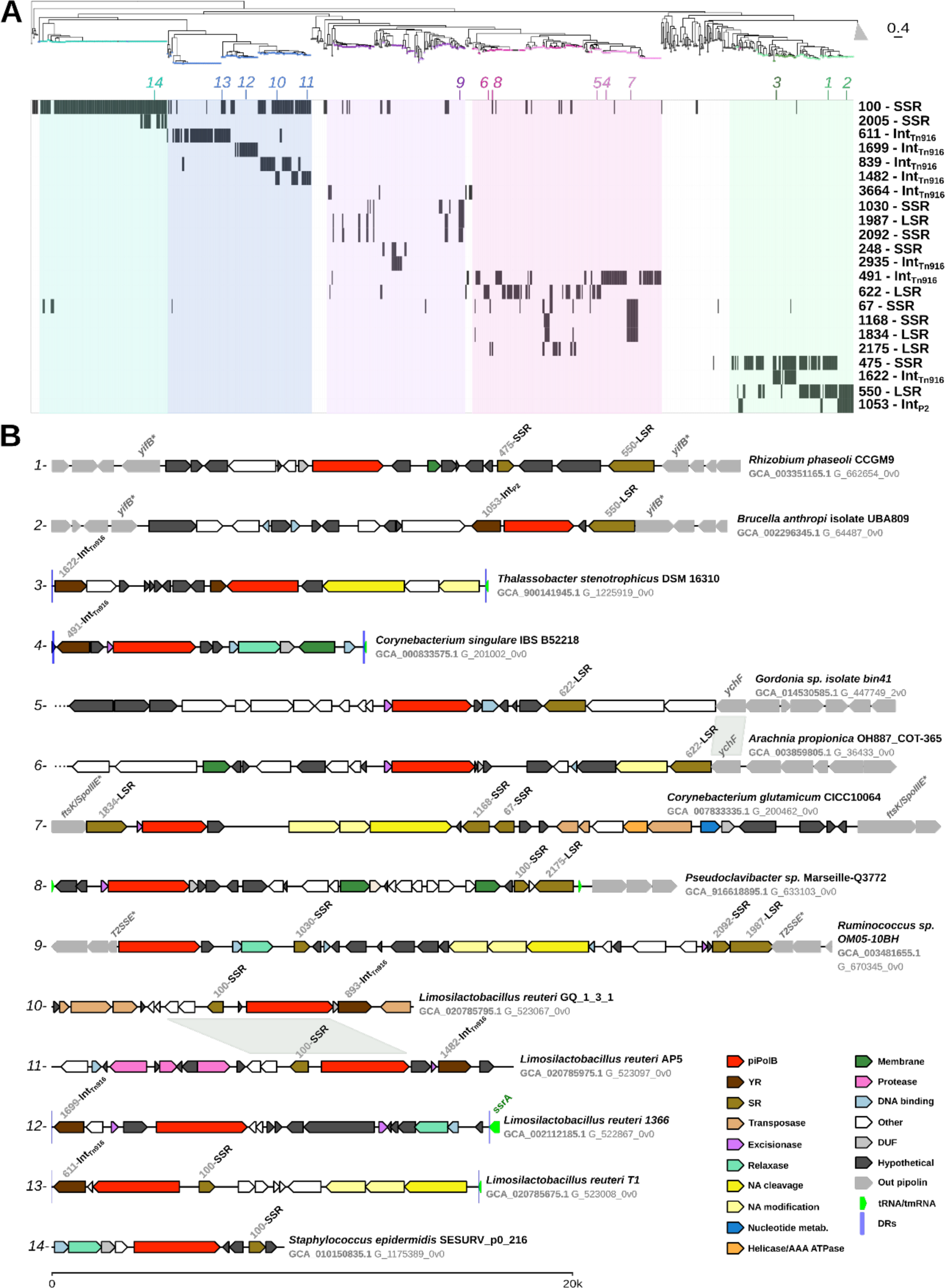
Pipolin recombinases outside *Gammaproteobacteria*. **A**. Maximum-likelihood phylogenetic tree show in Figure 3A with *Gammaproteobacteria* clade collapsed (grey triangle) to improve visualization of other clades. The presence-absence heatmap under the tree includes the candidate recombinases detected for each piPolB cluster (see methods). Italic numbers over the heatmap indicated the location of the pipolins represented below. **B**. Genomic organization of representative pipolins exemplifying the integrase exchanges and new integration sites detected in this study. Pipolin genes and other sequence features are represented as in Figure 3B, following the coloring scheme indicated in the legend. The initial three dots indicate unclear pipolin boundaries. Tyrosine and serine recombinases and their integration sites are labeled following their annotation and marked with an asterisk if they are truncated. Pipolins are also labeled indicating strain name, genome accession number and pipolin identifier.

In *Bacillota* (formerly *Firmicutes*), identification of for piPolB-associated integrases in *Clostridia* did not yield a clear pattern. Among a variety of diverse and low-frequency integrases, the most common is the association of a LSR is usually followed by a resolvase-like SRs (**Figure 4***, pipolin 9*). These recombinases are likely responsible for the integration of the element inside a T2SS E gene, leaving two truncated ORFs flanking the pipolin. On the contrary, integrases present in pipolins hosted in *Bacilli* class varied drastically depending on the piPolB subclade. Our results show that piPolBs in *Lactobacillaceae* are always associated to an YR of the Int_Tn916_ family (**Figure 4***, pipolins 10 to 13*) that show certain correlation with the piPolB phylogeny. Despite the prevalence of YRs we only found the integration site in pipolins encoding some groups of Int_Tn916_, either in a tRNA gene or in another novel site, the *ssrA* tmRNA gene, already known as integration site for MGEs in other diverse bacteria^62^. The other three Int_Tn916_ clusters associated with piPolB are usually encoded near a contig end so their integration site and DRs could not be detected. Also, like *Actinobacteria* pipolins, some *Lactobacillaceae* pipolin encode a small SRs besides the Int_916_ similar to resolvases and other SRs observed in pipolins from other gram-positive bacteria. In any case, these results indicate that pipolins in *Lactobacillaceae* are predominantly integrative elements while circular plasmid-like pipolins lacking integrases like pLME300 from *Lactobacillus fermentum* would be a rare exception. Regarding piPolBs from *Staphylococcus*, the other big clade of pipolins within the *Bacilli* group, no YR or LSR was found associated with piPolBs beyond plasmid resolvases. We thus hypothesize that pipolins in staphylococcus dwell in the bacteria as circular plasmids like pSE-12228-03 and pTnSha2^63^ (**Figure 4**, *pipolin 14*, **Figure S2, Figure S3**). Intriguingly, piPolBs from distant branches of the staphylococcal clade exhibit approximately 30% sequence identity, similar to mobilization genes, yet their genomic organization remains largely unchanged in several *Staphylococcus* species. This suggests that plasmid-pipolins in *Staphylococcus* are ancient and stable mobile genetic elements (MGEs), rather than a consequence of recent integrase loss.

Altogether, the newly identified integration sites allowed us to establish new boundaries for 761 pipolins. Combined with 134 *Staphylococcus* plasmids and 8,818 pipolins with DRs+tRNA/tmRNA, this results in a total of 9,713 (82.9 %) delimited pipolins. The diverse lifestyles of pipolins across different bacterial groups indicate that they have undergone significant structural changes during evolution, such as the exchange of an LSR for an Int_Tn916_. This highlights the piPolB gene as the sole hallmark of these elements, while enabling a variety of integration mechanisms and lifestyles by utilizing additional factors likely exchanged from other mobile genetic elements.

### 3. Diverse defense genes constitute the cargo of pipolins

With the aim of characterizing the genetic composition of pipolins we used specialized tools to annotate phage (PHROGs^45^) and conjugation (CONJScan^46^) genes, defense systems (PADLOC^49^), virulence factors (VFDB^50^), and ARGs (AMRFinderPlus^48^). Then, in order to compare the pipolin genetic composition with the main MGE groups, the same methods were applied to a plasmid database (PLSDB^41^), a comprehensive bacterial virus dataset (sequences from RefSeq, Genbank, EMBL and DDBJ contained in PhageScope^44^), and a conjugative and integrative MGEs (ciMGEs) database. After removing redundant sequences and restricting by size (see Methods), we obtained a full dataset consisting of 7,409 pipolins, 50,022 plasmids, 7,050 phages, and 10,925 ciMGEs **(Supplementary Dataset 4).**

PHROGs profiles allowed us to annotate phage functions in almost 10% of pipolin genes, the lowest value of the four MGE groups (**Figure 5A**). However, phage-related genes in all groups of pipolins mainly belong to typical pipolin functions, such as “integration and excision”, “DNA, RNA and nucleotide metabolism”, and “other” classes, while genes involved in phage structure or lysis are practically absent compared with the other MGE groups (**Figure S4**).

**Figure 5.**
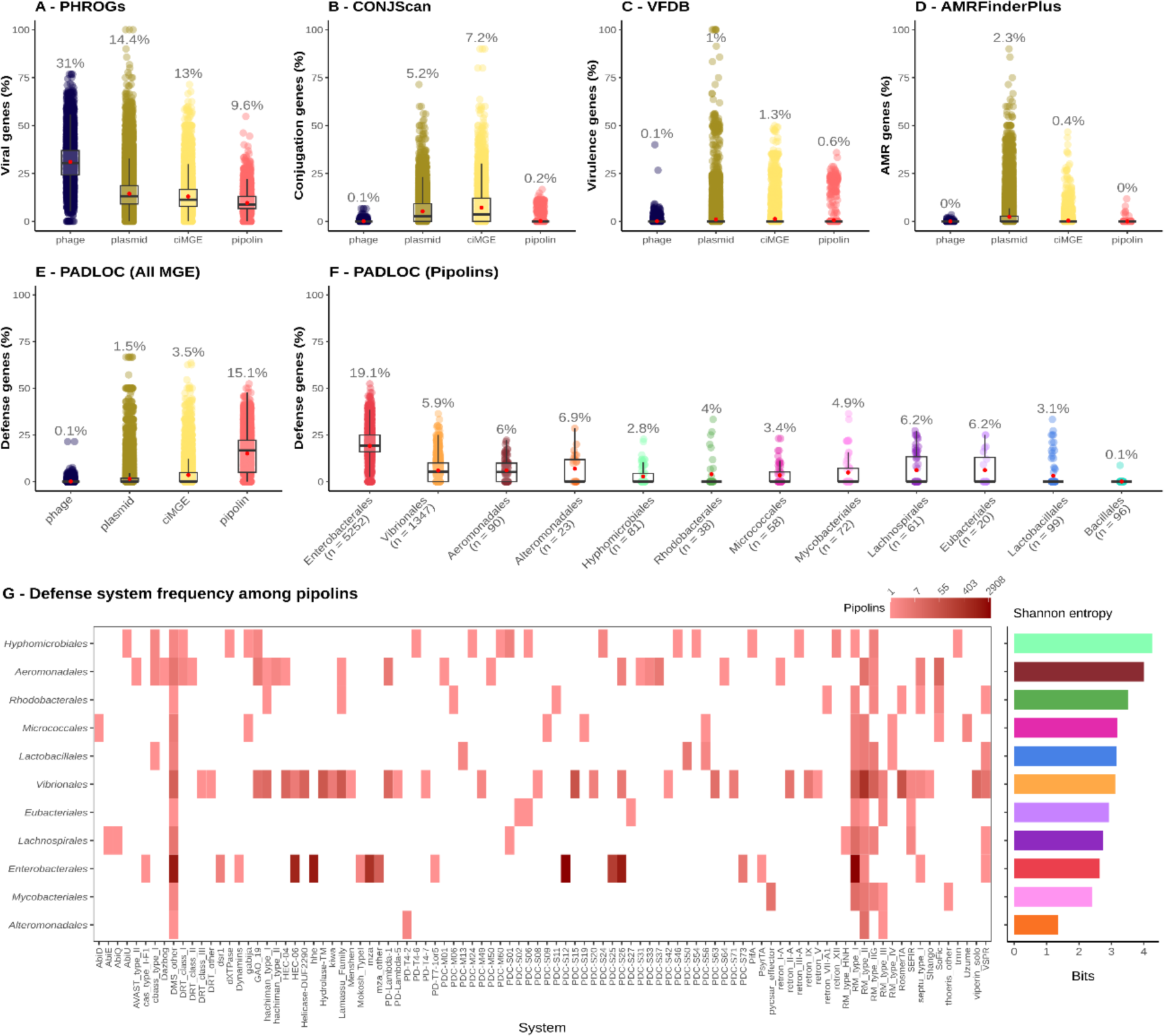
Comparative annotation of pipolins and other MGEs. **A-E**. Boxplots indicating the frequency of genes annotated by (**A**) PHROGs profiles, (**B**) CONJScan profiles, (**C**) VFDB sequence search, (**D**) AMRFinderPlus, and PADLOC (**E**) for each MGE category. Individual MGE values are represented by dots in scatterplots behind the boxplots. Red points in each category represent the average value, whose numerical value appears above the scatterplot. **F**. PADLOC annotation result in pipolins, grouped by host order. Numbers below order name indicate sample size (*i.e.* number of pipolins in that order). Only n >= 20 are represented. **G.** Heatmap showing which defense systems are found in pipolins of the most frequent orders. Adjacent barplot represents the diversity of systems found in each order measured by the Shannon entropy. The entropy for each order is defined by *H_order_* = − ∑ *p_s_* log(*p_s_*), where *p_s_* is the observed proportion of pipolins containing the system *s*. This is equivalent to the observed proportion of systems *s* encoded in pipolins from that order if we assume that a pipolin does not contain more than one system of the same class (for instance, two Type I RM systems).

Mobilization genes could be detected in ciMGEs and plasmids but were rare in phages and especially in pipolins (**Figure 5B**). Only pipolins from gram-positive bacteria encode MOBP or MOBV relaxases, the latter usually along with a VirB11 ATPase (**Figure S4**). Full or partial T4SSs were not found in any pipolin, congruent with the relatively small size of these elements. Thus, relaxase-encoding pipolins groups could be considered as piPolB-encoding Integrative and Mobilizable Elements (IME-pipolins), except for staphylococcal pipolins where there is no YR/LSR and could be simply defined as piPolB-encoding mobilizable plasmids (plasmid-pipolins).

Besides the lack of phage or plasmid structural genes, the annotation of cargo genes in pipolins also revealed striking differences. While plasmids and ciMGEs encoded a variety of known AMRs or virulence genes, they are very uncommon in phages, in line with previous studies^51^ (**Figure 5CD**). Regarding pipolins, these MGEs are devoid of known antimicrobial and virulence genes, except for a group of nearly 200 *E. coli* pipolins that encode a gene cluster related to an exopolysaccharide export system, which would be a recent acquisition as it had not yet been found in other enterobacteria. Lastly, defense functions annotated with PADLOC represented 15.9% of pipolin genes (**Figure 5E-F**), several times superior that the average for both integrated and circular elements. This result places pipolins as one of the MGE families with the highest density of defense genes, qualifying them as defense islands. Pipolins from *Gammaproteobacteria* contributes the most to the pool of defense genes in pipolins, which are one out of five genes in *Enterobacterales*. The remaining groups contain roughly 5% of defense genes, except *Bacillales*, where defense genes are extremely rare. In total, up to 98 different system classes were detected (excluding the “DMS_other’’ class). The orders with the greatest diversity of systems are *Vibrionales*, *Aeromonadales,* and *Hyphomicrobiales,* which showed the highest entropy value (**Figure 5G**). Despite being the largest group by far, both in terms of elements and defense genes, *Enterobacterales* pipolins rank fourth in number of different systems and ninth (of 11 orders) in entropy, which may be explained by the late arrival of pipolins to this order.

These results, along with the overall scarcity of structural genes, indicate that pipolins are specialized to accommodate a variety of cargo genes, with a strong bias towards defense systems, especially in enterobacterial pipolins, whereas AMR genes and virulence factors are reluctant to colonize pipolins.

### 4. High variability of pipolins explained by the detection of recombining genes

#### 4.1 Pipolins show preferent recombination with plasmids and ciMGEs than phages

The diversity of accessory or cargo genes encoded by pipolins, well represented by their repertoire of defense genes, suggest that pipolins are active elements that interact and exchange genes with other MGEs. To address whether pipolins contribute to genetic exchange between MGEs, we adapted a recent approach to detect recent gene exchanges based on the weighted gene repertoire relatedness (wGRR) shared by the elements^51^. As detailed in Methods, homologous recombining genes (RGs) between two MGEs can be detected if their sequence identity and coverage is similar enough (>80% aminoacid identity and >80% coverage of both sequences) and are encoded in quite different MGEs (*i.e.* element pairs with low wGRR). Otherwise, they are classified as non-recombining genes (NRGs). Elements that had no highly similar genes in pipolins or only shared ubiquitous ISs were discarded. The resultant dataset included 4,239 plasmids, 412 ciGMEs, and 17 phages, apart from the initial 7,409 pipolins. This makes up 8.47% of plasmids, 5.84% of ciMGEs, and less than 0.2% of phages from the original dataset, hinting that pipolins are more prone to exchange genetic material with conjugative elements than phages.

Best-bidirectional hits (BBHs) with >80% aminoacid identity and >80% coverage calculated for our data show a bimodal distribution of their associated wGRR values (*i.e.* the wGRR of the elements encoding the pair of homologues), where the lower heap of wGRR values is compatible with exchange events according to previous studies^64^ (**Figure S5**). We set a threshold of wGRR < 0.2 to obtain a percentage of pipolin genes classified as RG was 40%, with a mean RG/NRG ratio of 0.41, although highly group-dependent (**Figure 6A-B**). In *Vibrionales* and *Enterobacterales* around half of the genes are considered RGs while in the other of groups the RGs are below 20%, correlating with the group size. Pipolins in average encode less RGs than plasmids but more than ciMGEs, which is in line with previous reports showing that conjugative plasmids are more variable and prone to genetic exchange than integrated elements like ICEs^64^. In fact, in our pipolin-oriented dataset, most of the recombination events were detected in plasmid-to-plasmid exchanges. Regarding pipolins, recombination between pipolins and plasmids are the most frequent event comprising 47.9% of gene families involved in exchanges while pipolin-pipolin and pipolin-ciMGEs contributed with 27.7% and 23.4%, respectively (**Figure 6C**). A graph containing all MGEs linked by shared RG confirms that genetic exchange between pipolins and plasmids and ciMGEs takes place in pipolins from all phyla (**Figure 6D**).

**Figure 6.**
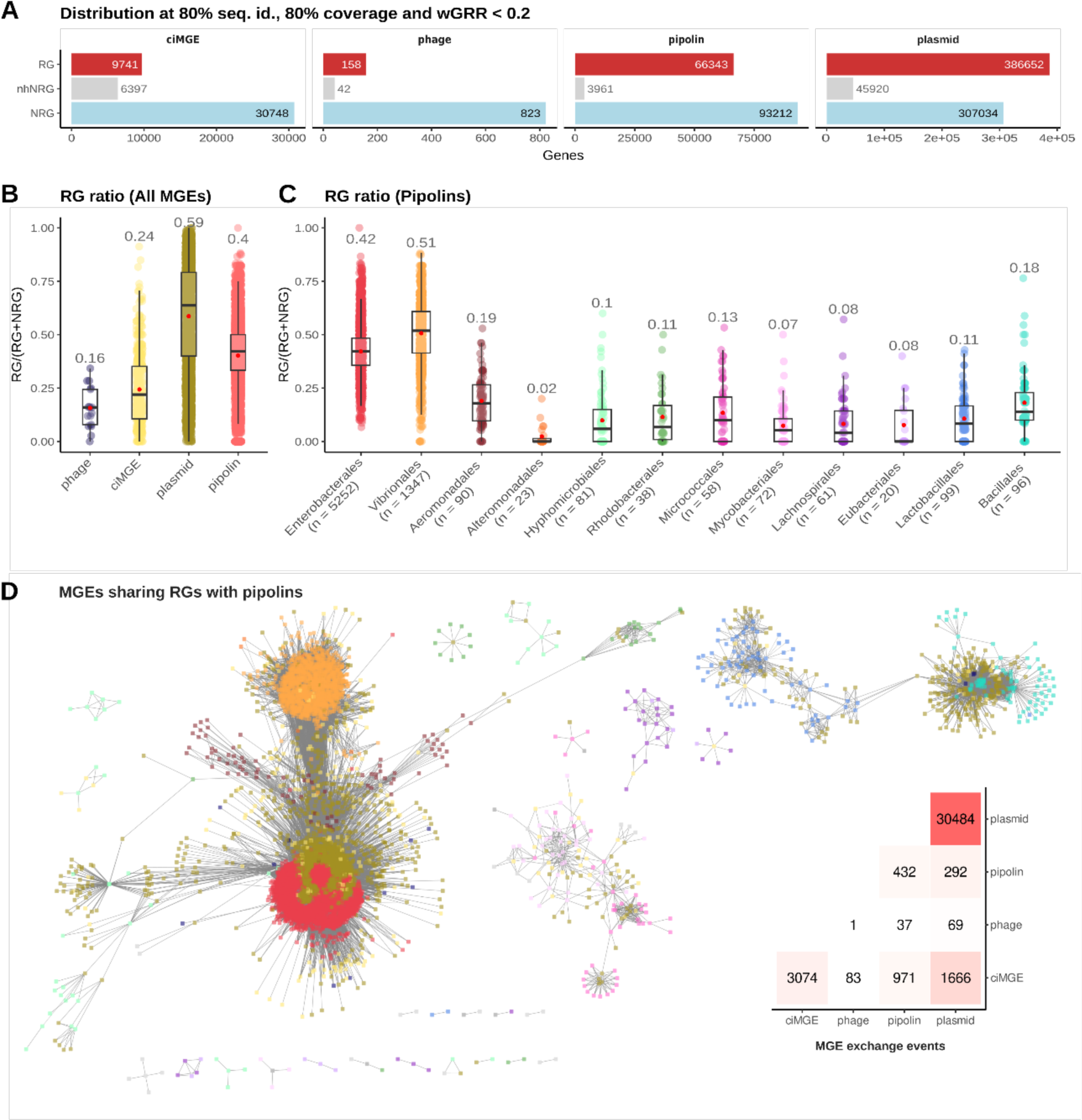
RG computation in pipolins and other MGEs. **A**. Counts of RG, NRG, and nhNRG for each MGE class after setting wGRR < 0.2, 80% identity, and 80% coverage as threshold for RG identification. **B.** Boxplots showing the proportion of RGs, given by the RG/(RG+NRG) ration, in each MGE group. Individual MGE values are represented as dots in scatterplots behind the boxplots. Red points in each category represent the average value, whose numerical value is shown above the scatterplot. **C**. RG frequency in pipolins, grouped by host order. Numbers below order name indicate sample size (*i.e.* number of pipolins in that order). Only n >= 20 are represented. **D.** Graph showing genetic exchange events involving pipolins. Nodes represent MGEs, which are linked by an edge if they share an RG was detected between them. Plasmids, ciGMEs, and phages are colored as in D and pipolins are colored as in E. Graph plotting was carried out in Cytoscape 3.10 under a prefuse force directed layout (default parameters). Heatmap on the bottom right side of the graph shows the counts of RG family exchange events between MGE classes (see Methods).

These results show that pipolins are actively involved in genetic exchange processes that take place in bacterial populations. In groups with large sample size (*i.e. Enterobacterales* and *Vibrionales*) the ratio of RGs is closer to plasmids than ciMGEs or phages, which indicates that pipolins may be characterized by a higher flexibility than their integrated counterparts.

#### 4.2. Pipolins defense genes undergo extensive exchange with other GMEs in Enterobacteria

In order to disclose the functions of pipolin genes involved in recent exchanges, we assessed for each functional category whether there is a statistically significant overrepresentation of RGs (see Methods) in each group of pipolins (**Supplementary Data 5**). In pipolins from *Enterobacterales*, there is an enrichment of RGs in genes related to nucleic acid metabolism, auxiliary functions, and other functions but not in genes involved in integration and excision. Indeed, we detected an enrichment of the most frequent defense systems (except PDC-S26), which have been recently exchanged with other plasmids and ciMGEs where the systems are usually found near transposases (**Figure 7**). Furthermore, these defense gene exchanges involve both frequent^20^ (Type 1 RM, PDC-S12) and rare systems (mza, hhe, HEC-06), which show a patchy presence-absence pattern suggestive of recombination events (**Figure S6**). Virulence genes related to immune modulation were significantly enriched as they have also been recently exchanged with a conjugative element. In contrast, pipolin core genes in this group, such as the YRs, WYL and DUF2787 containing genes are mostly classified as NRGs (**Figure S7**) as they lack a homologue in any distant MGE. The piPolB genes, however, are mainly classified as RGs since there is a group of pipolins with a deletion of the core genes, adjacent to a cointegrated phage that shares low wGRR values with the rest of enterobacterial pipolins (**Figure 3***, pipolin 2*). Thus, pipolins in *Enterobacteria* are bimodular, with a conserved genetic core near the integration site, while the distal site can serve as a platform for different cargo genes that can be easily exchangeable with other MGEs, which would favor fast host adaptation.

**Figure 7.**
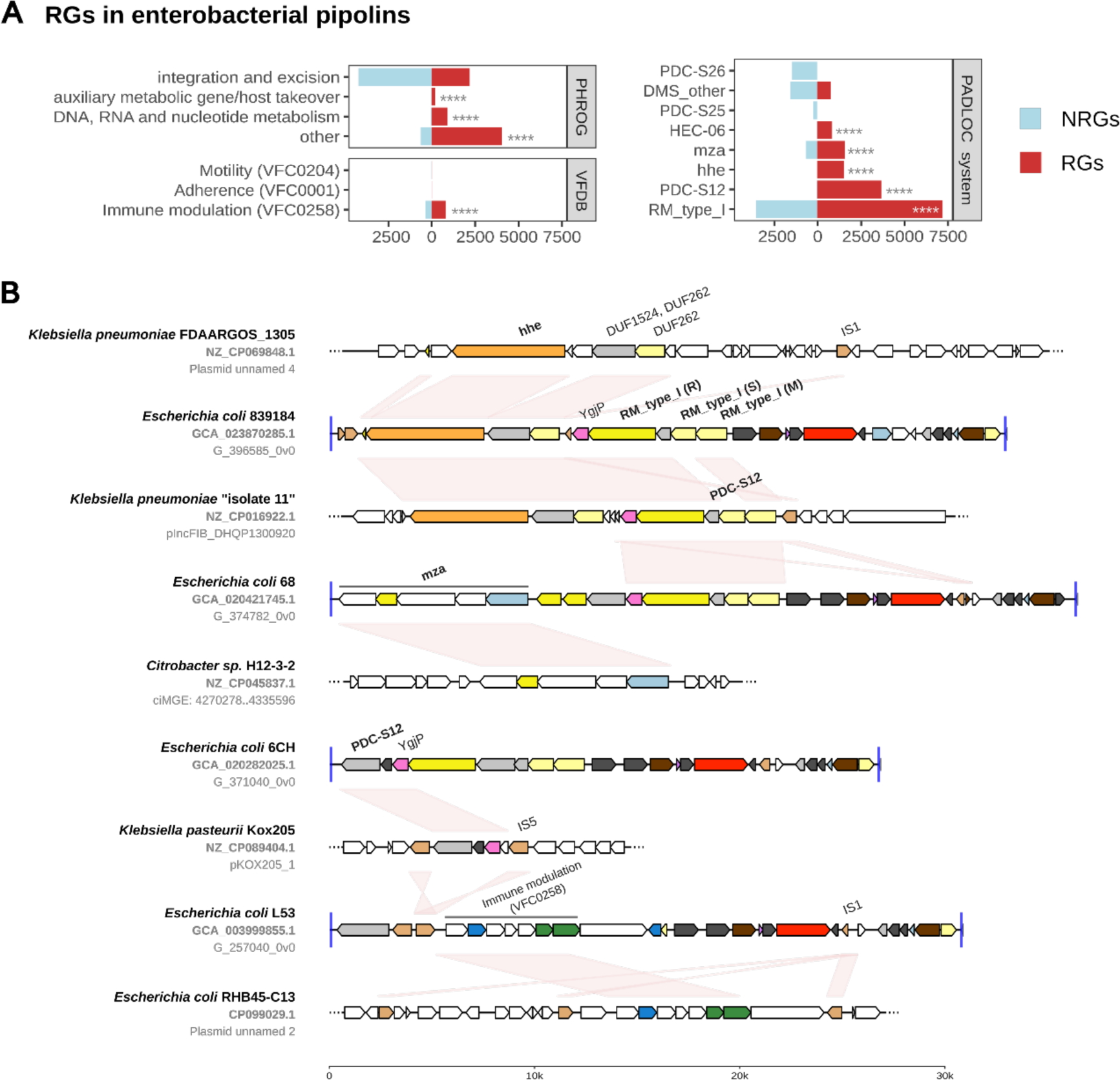
Exchange of cargo genes between pipolins and conjugative elements in *Entetrobacterales*. **A.** Number of RGs and NRGs detected in enterobacterial pipolins grouped by each category of annotations (PHROG categories, VF classes, and PADLOC systems). The RG count of each category was compared to the number of NRGs using a Fischer’s exact test with the Benjamini-Hochberg multiple testing correction. Significant overrepresentation of RGs of a category is shown as: * <= 0.05, ** <= 0.01, *** <= 0.001, **** <= 0.0001. Statistical overrepresentation of NRGs and the RG/NRG count of unannotated genes is not shown in the plot, but data and test results can be consulted in Supplementary Data 10. **B.** Genomic organization of representative pipolins exemplifying cargo exchanges with other MGEs. Genes and other sequence features are represented as in Figure 3 and 4, following the coloring scheme indicated in the legends. Due to the size difference between pipolins and the other MGEs, only the recombining region of plasmids and ciMGEs is shown (Three dots indicate that sequence continues in that direction). MGEs are labeled indicating strain name, genome or plasmid accession number and element identifier or plasmid name. Red links mark RGs between sequences. RG annotation is displayed above a representative gene and marked in bold if corresponds to a PADLOC defense system.

#### 4.3. Integrative pipolins show a genetic plasticity characteristic of extrachromosomal MGEs

In pipolins from the other gammaproteobacterial orders the RGs mainly belong to the core pipolin functions instead of the accessory variable functions (**Figure 8A-B**). Although pipolins from *Vibrionales* and *Aeromonadales* show the highest system diversity, only three systems were significantly enriched in RGs: SoFic and PDC-37 in *Aeromonadales*, RosmerTA in *Vibrionales*. In contrast, recurrent pipolin genes annotated as integrases, DUF2787, WYL and the own piPolB comprise most of the RG set in these taxa (**Figure S7**). Strikingly, we found the integrases to be exchanged with many plasmids and ciMGEs but the remaining RG-core genes were found exclusively on pipolins, with the exception of 2 ciMGEs cointegrated next to a pipolin. Integrase exchange would explain the target tRNA changes observed in certain *Vibrionales* and *Aeromonadales* subgroups (**Figure 8B** – pipolins 1 to 4, **Figure S1**), as different Int_SXT_ subfamilies may show preferential insertion over different tRNA genes^40,65^. Alternatively, due to the reduced size of the pipolin genetic core, fast genetic exchange of cargo genes would result in pipolins with very different genetic repertoires, but with high sequence similarity between core genes that are classified as RGs. Therefore, these results are consistent with both exchange of core gene between pipolins and rapid exchange of cargo genes over short evolutionary time scales. Moreover, we found a similar result in several groups beyond *Gammaproteobacteria*, where we also found integrase exchange events that explain the incongruencies between the piPolB phylogeny and the presence-absence pattern observed in this study.

**Figure 8.**
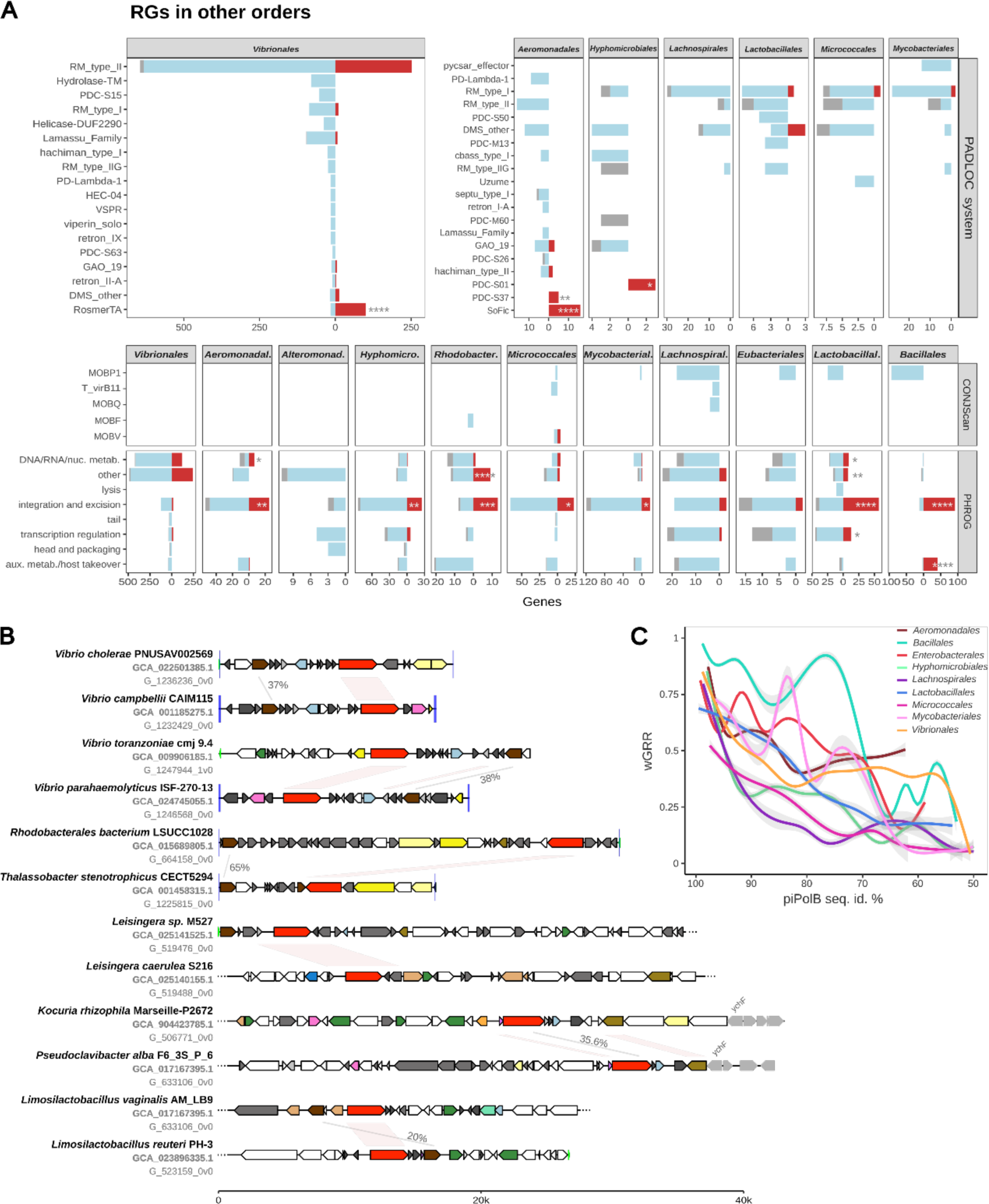
Pipolin core gene exchange. **A.** Number of RGs and NRGs detected in pipolins grouped by each category of annotations (PHROG categories, VF classes, and PADLOC systems) and bacterial order. The RG count of each category was compared to the number of NRGs using a Fischer’s exact test with the Benjamini-Hochberg multiple testing correction. Significant overrepresentation of RGs of a category is shown as: * <= 0.05, ** <= 0.01, *** <= 0.001, **** <= 0.0001. Statistical overrepresentation of NRGs and the RG/NRG count of unannotated genes is shown in Supplementary Data 10. **B.** Genomic organization of representative pipolins exemplifying integrase and piPolB exchanges with other pipolins. Genes and other sequence features are represented as in Figure 3 and 4. Pipolins with unclear boundaries are trimmed (three dots in the plot) for representation purposes. Red links mark RGs between sequences. To remark contrast with RGs, sequence identity for some piPolBs and integrases linked by grey lines is shown. **C.** Variation of wGRR values as a function of the piPolB sequence identity. Smoothed curves were calculated for each order (only orders with >50 pipolins) using generalized additive models (GAM^66^).

We also identified as significantly enriched RGs the abovementioned IS1272-family transposase and a NADH-dependent trans-2-enoyl-acyl carrier protein reductase (*fabI*), which conform a transposable element named TnSha1 since *fabI* derives from a copy of the *fabI* from *Staphylococcus haemolyticus* that grants resistance to antimicrobials from the triclosan family^63^. This transposable element has been exchanged between multiple plasmids and ciMGEs and, although restricted to *Staphylococcus* plasmid-pipolins, it is the only clear association between pipolins and antimicrobial genes to date. Interestingly, we found alternative versions lacking the IS1272, leaving *fabI* in the plasmid, and plasmid co-integrates encoding other AMR genes, plasmid replication and partition proteins, but at the cost of disrupting the piPolB gene (**Figure S3)**.

Finally, we quantified the pipolin rate of change by modelling the wGRR value as a function of the piPolB divergence (*i.e.* the aminoacid sequence identity) similarly to previous experiments with MOB_P_ and MOB_F_ plasmids^53^. The resultant functions inferred by the data showed that wGRR values between pipolins decay with rates similar to MOB_P_ and MOB_F_ plasmids (**Figure 8D**), further supporting the idea that pipolins show flexibility levels nearer to plasmids than integrated MGEs.

In conclusion, our results showed that pipolins are active members of the MGE genetic exchange network in bacteria populations. This capacity is well exemplified by the RGs detected in enterobacterial pipolins, which are mainly defense-related cargo genes found in a variety of plasmids and ciMGEs. Overall, we conclude that pipolins are flexible short-sized elements that renew their cargo content in rates similar to extrachromosomal MGEs.

## CONCLUDING REMARKS

We report here the presence of pipolins in 11,431 of more than 1.3 million bacterial genome assemblies, spanning diverse taxa from major bacterial clades, including unforeseen prevalence in hosts of biomedical and biotechnological interest such as *Vibrio*, *Aeromonas*, and *Limosilactobacillus,* where approximately 10% of genomes contain pipolins. The observation that several genomes of these taxa contain more than one pipolin and have low piPolB identity suggests that pipolins have been present for a long time compared to other groups such as Enterobacterales. This idea is supported by both the phylogenetic analysis of piPolB and the broader genetic core observed in other Gammaproteobacteria groups. In fact, previous analysis of *E. coli* defense hotspots found the pipolin insertion site (tRNA-Leu between *psgA* and *yecA*) to be occupied in 21% of genomes, mainly by prophages (80%) and unidentified MGEs (20%)^67^, explaining the lower prevalence observed in this species. Despite the later arrival of pipolins in the Enterobacteriales, these elements have successfully established themselves in all major genera and show evidence of a recent HGT between them. This confirms that pipolins are currently active MGEs capable of transferring to other enterobacteria that do not possess a known conjugation or viral gene beyond integrase.

Analysis of pipolin encoded integrases combined with the phylogenetic analysis of the piPolB further confirmed that pipolins in *Enterobacteriales* derives from a common gammaproteobacteria ancestor that would contain a Int_P2_ followed by an excisionase, a Int_SXT_, and DUF2787 and WYL domain containing proteins, suggest a common evolutionary origin and functional association between the piPolB and these genes. These core genes are reliable phylogenetic markers of pipolins in this group, however, in other groups we found several integrase exchange events that highlighted pipolin versatility. Furthermore, these recombination events would explain how pipolins have colonized new alternative sites, sometimes different than a tRNA (*yifB*, *ychF*, *FtsK/SpoIII*, *T2SS*). Despite that plasticity, most pipolins encode either a YR or an LSR, highlighting pipolins as an MGE family of integrative elements. The only group of pipolins lacking YR or LSR are found in *Staphylococcus*, where pipolins are circular plasmids similar to pSE-12228-03 and pTnSha2^63^.

Pipolins carry a sizable proportion of cargo genes with a minimal core of conserved genes. However, they lack classic adaptive traits common in other MGEs, such as AMR genes and virulence factors, which have only recently been acquired in a few specific cases. In contrast, they show a strong preference for defense systems, and devote a larger proportion of their gene repertoire to them than phages, plasmids, and ciMGEs. This observation is in line with previous results reporting an inverse correlation between the presence of defense systems and the presence of AMR and virulence genes^42^. Gathering of defense genes in close proximity is well documented^68–70^ and, at least in *E. coli*, that accumulation can be favored by the presence of a variety MGEs in specific location of chromosome hotspots, including tRNA pipolin integration sites^67^, providing synergistic defense capacity^71^. However, the gene flux rate of defense systems and their interaction with different MGEs is unclear. The evolution of defense systems is driven by the bacteria-phage arm race and can be very fast^70^, but it is also determined by fitness cost, autoimmunity and HGT barriers^11,72^. The identification of recombining genes between pipolins and plasmids and ciMGEs revealed a high exchange rate of pipolin cargo genes, which include defense genes (**Figure 7A** and **Figure 8A**) and a variety of proteins containing DUFs or domains involved in nucleic acids metabolism that might also be involved in defense (such as nucleases, helicases or DNA glycosylases^69,73,74^) or even in defense evasion^75^ (**Figure S7**).

All in all, our results support a model in which pipolins are bimodular defense islands, with a minimalist genetic core responsible for the maintenance of the element and a second dynamic module dedicated to building a highly variable defensive arsenal in frequent exchange within the bacterial mobilome. In this respect, pipolins resemble other recently reported integrative defense-related elements such as *Pseudomonas aeruginosa* cDHS^76^ or GMT islands^77^ that, like pipolins, are dedicated to maintaining a reservoir of defense factors together with a reduced set of core genes. We propose that these groups of GMEs, as well as other previously overlooked elements, may represent a novel superfamily of GMEs that would serve as a platform with a dynamic catalog of defense systems, which will be eventually incorporated by ciMGEs, plasmids, and other elements endowed with gene transfer machinery. Thus, the autonomous elements would benefit from an orthogonal and dynamic reservoir of defense genes provided by pipolins and other defense elements lacking their own mobilization machinery, recently referred to as hitchhikers^78^, less constrained by the limitations of HGT by defense systems in the short term^79^. In exchange, non-sensitive helper elements would eventually provide the means for their eventual mobilization. A comprehensive analysis of the gene exchange rates of defense factors in distinct types of MGEs may shed light on the dynamics of defense systems between autonomous and defense-hitcher elements.

## ACKNOLEDGMENTS

This work was funded by MCIN/AEI/10.13039/501100011033 and ERDF A way of making Europe [PID2021-123403NB-I00] and Comunidad de Madrid (V PRICIT call Research Grants for Young Researchers from Universidad Autónoma de Madrid) [SI3/PJI/2021-00271]. VMC was holder of an FPI-UAM PhD Fellowship from UAM [SFPI/2023-00603].

We would like to thank Liubov Chuprikova for helping us with ExplorePipolin technical issues and providing insights regarding pipolin structure. Additionally, we give special thanks to Mario Rodríguez-Mestre for his contributions on gene annotation and clustering methods, particularly with the MMSeqs2 usage.

## DATA AVAILABILITY

The genomics data employed in this study is openly accessible and can be obtained from the NCBI databases (https://www.ncbi.nlm.nih.gov/) using the respective accession IDs. Large datasets, spanning pipolins annotation and representation, full list of GME genes prediction and functional annotation and details about recombining genes (RG) detection have been deposited in e-Cienciadatos repository with DOI: <u>10.21950/QG3QEE</u>

## CODE AVAILABILITY

Custom scripts (pipolin screening and analysis, wGRR calculation, quantification of gene flow and enrichment tests) used in this study are available in GitHub: https://github.com/rnrlab/pipolin_bacteria_screening

