## Supplementary Information for "Pipolins are bimodular platforms that maintain a reservoir of defense systems exchangeable with various bacterial genetic mobile elements"

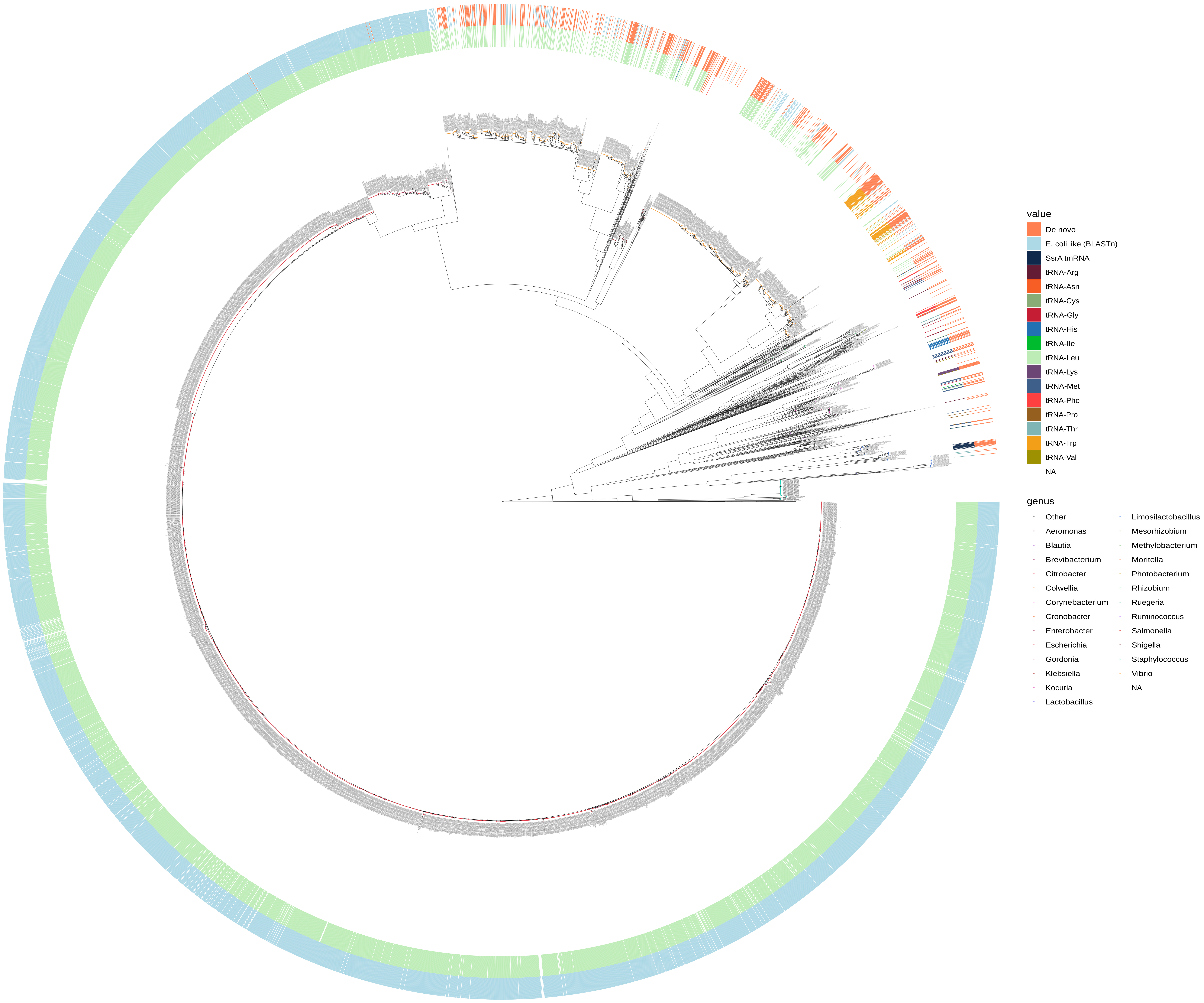
value
De novo
E. coli like (BLASTn)
SsrA tmRNA
tRNA-Arg
tRNA-Asn
tRNA-Cys
tRNA-Gly
tRNA-His
tRNA-Ile
tRNA-Leu
tRNA-Lys
tRNA-Met
tRNA-Phe
tRNA-Pro
tRNA-Thr
tRNA-Trp
tRNA-Val
NA
genus
Other
Aeromonas
Blautia
Brevibacterium
Citrobacter
Colwellia
Corynebacterium
Cronobacter
Enterobacter
Escherichia
Gordonia
Klebsiella
Kocuria
Lactobacillus
Limosilactobacillus
Mesorhizobium
Methylobacterium
Moritella
Photobacterium
Rhizobium
Ruegeria
Ruminococcus
Salmonella
Shigella
Staphylococcus
Vibrio
NA

**Figure S1. Phylogenetic inference of the piPolB gene based on the protein sequence and integration site**. The phylogeny displayed corresponds with a maximum-likelihood phylogenetic tree of 9,740 piPolBs and Bam35 DNAP (outgroup) inferred by IQ-TREE v2 (see Methods). ModelFinder best-fit model was Q.pfam+I+I+R10 according to BIC. Colored tips represent the most frequent genera, following the legend at the right. The inner ring surrounding the tree indicates the integration site of the pipolin encoding the piPolB in the tip and the outer ring indicates the type of direct repetitions, as detected by ExplorePipolin. Numbers next to integrase type corresponds to cluster number. Tree tips are labeled with piPolB and pipolin identifier, genome assembly accession number and species name (if available), in that order. Numbers over the tree branches indicates bootstrap values.

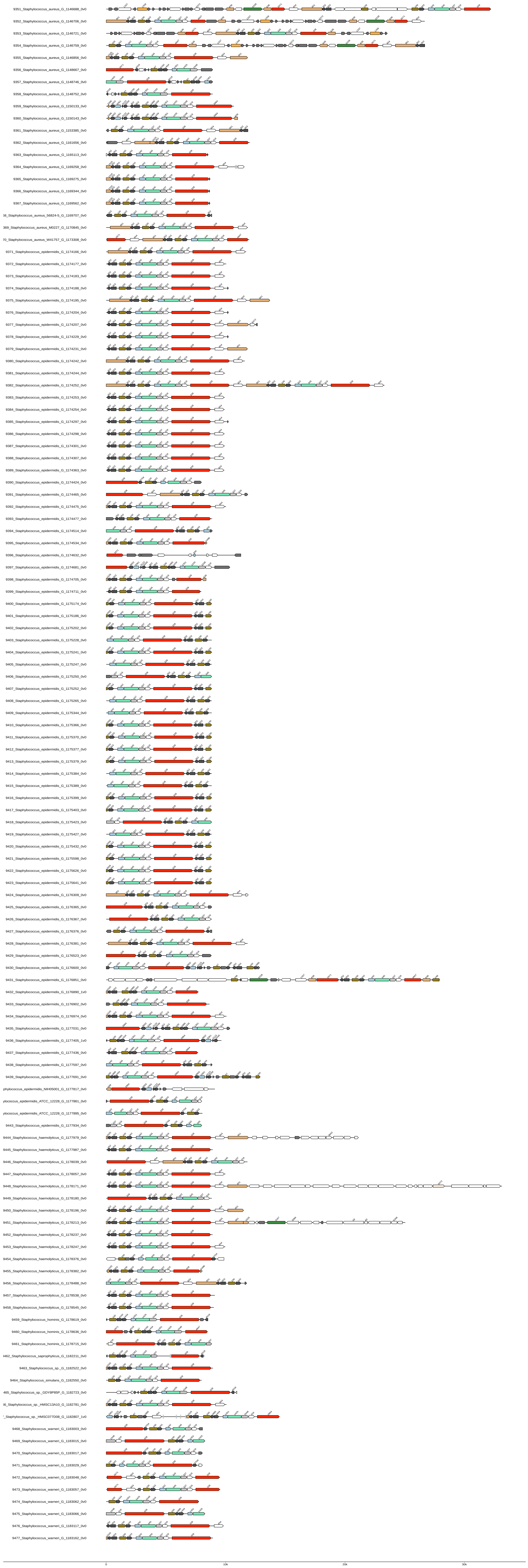

9351_Staphylococcus_aureus_G_1146688_0v0
9352_Staphylococcus_aureus_G_1146706_0v0
9353_Staphylococcus_aureus_G_1146721_0v0
9354_Staphylococcus_aureus_G_1146759_0v0
9355_Staphylococcus_aureus_G_1146856_0v0
9356_Staphylococcus_aureus_G_1148667_0v0
9357_Staphylococcus_aureus_G_1148746_0v0
9358_Staphylococcus_aureus_G_1148752_0v0
9359_Staphylococcus_aureus_G_1150133_0v0
9360_Staphylococcus_aureus_G_1150143_0v0
9361_Staphylococcus_aureus_G_1153385_0v0
9362_Staphylococcus_aureus_G_1161656_0v0
9363_Staphylococcus_aureus_G_1165113_0v0
9364_Staphylococcus_aureus_G_1169258_0v0
9365_Staphylococcus_aureus_G_1169275_0v0
9366_Staphylococcus_aureus_G_1169344_0v0
9367_Staphylococcus_aureus_G_1169562_0v0
68_Staphylococcus_aureus_56824-5_G_1169707_0v0
369_Staphylococcus_aureus_M0227_G_1170845_0v0
70_Staphylococcus_aureus_W41757_G_1173308_0v0
9371_Staphylococcus_epidermidis_G_1174166_0v0
9372_Staphylococcus_epidermidis_G_1174177_0v0
9373_Staphylococcus_epidermidis_G_1174183_0v0
9374_Staphylococcus_epidermidis_G_1174188_0v0
9375_Staphylococcus_epidermidis_G_1174195_0v0
9376_Staphylococcus_epidermidis_G_1174204_0v0
9377_Staphylococcus_epidermidis_G_1174207_0v0
9378_Staphylococcus_epidermidis_G_1174229_0v0
9379_Staphylococcus_epidermidis_G_1174231_0v0
9380_Staphylococcus_epidermidis_G_1174242_0v0
9381_Staphylococcus_epidermidis_G_1174244_0v0
9382_Staphylococcus_epidermidis_G_1174252_0v0
9383_Staphylococcus_epidermidis_G_1174253_0v0
9384_Staphylococcus_epidermidis_G_1174254_0v0
9385_Staphylococcus_epidermidis_G_1174297_0v0
9386_Staphylococcus_epidermidis_G_1174298_0v0
9387_Staphylococcus_epidermidis_G_1174301_0v0
9388_Staphylococcus_epidermidis_G_1174307_0v0
9389_Staphylococcus_epidermidis_G_1174363_0v0
9390_Staphylococcus_epidermidis_G_1174424_0v0
9391_Staphylococcus_epidermidis_G_1174465_0v0
9392_Staphylococcus_epidermidis_G_1174475_0v0
9393_Staphylococcus_epidermidis_G_1174477_0v0
9394_Staphylococcus_epidermidis_G_1174514_0v0
9395_Staphylococcus_epidermidis_G_1174534_0v0
9396_Staphylococcus_epidermidis_G_1174632_0v0
9397_Staphylococcus_epidermidis_G_1174681_0v0
9398_Staphylococcus_epidermidis_G_1174705_0v0
9399_Staphylococcus_epidermidis_G_1174711_0v0
9400_Staphylococcus_epidermidis_G_1175174_0v0
9401_Staphylococcus_epidermidis_G_1175186_0v0
9402_Staphylococcus_epidermidis_G_1175202_0v0
9403_Staphylococcus_epidermidis_G_1175228_0v0
9404_Staphylococcus_epidermidis_G_1175241_0v0
9405_Staphylococcus_epidermidis_G_1175247_0v0
9406_Staphylococcus_epidermidis_G_1175250_0v0
9407_Staphylococcus_epidermidis_G_1175252_0v0
9408_Staphylococcus_epidermidis_G_1175265_0v0
9409_Staphylococcus_epidermidis_G_1175344_0v0
9410_Staphylococcus_epidermidis_G_1175366_0v0
9411_Staphylococcus_epidermidis_G_1175370_0v0
9412_Staphylococcus_epidermidis_G_1175377_0v0
9413_Staphylococcus_epidermidis_G_1175379_0v0
9414_Staphylococcus_epidermidis_G_1175384_0v0
9415_Staphylococcus_epidermidis_G_1175389_0v0
9416_Staphylococcus_epidermidis_G_1175399_0v0
9417_Staphylococcus_epidermidis_G_1175403_0v0
9418_Staphylococcus_epidermidis_G_1175423_0v0
9419_Staphylococcus_epidermidis_G_1175427_0v0
9420_Staphylococcus_epidermidis_G_1175432_0v0
9421_Staphylococcus_epidermidis_G_1175598_0v0
9422_Staphylococcus_epidermidis_G_1175626_0v0
9423_Staphylococcus_epidermidis_G_1175641_0v0
9424_Staphylococcus_epidermidis_G_1176309_0v0
9425_Staphylococcus_epidermidis_G_1176365_0v0
9426_Staphylococcus_epidermidis_G_1176367_0v0
9427_Staphylococcus_epidermidis_G_1176376_0v0
9428_Staphylococcus_epidermidis_G_1176381_0v0
9429_Staphylococcus_epidermidis_G_1176523_0v0
9430_Staphylococcus_epidermidis_G_1176600_0v0
9431_Staphylococcus_epidermidis_G_1176851_0v0
9432_Staphylococcus_epidermidis_G_1176890_1v0
9433_Staphylococcus_epidermidis_G_1176902_0v0
9434_Staphylococcus_epidermidis_G_1176974_0v0
9435_Staphylococcus_epidermidis_G_1177031_0v0
9436_Staphylococcus_epidermidis_G_1177405_1v0
9437_Staphylococcus_epidermidis_G_1177436_0v0
9438_Staphylococcus_epidermidis_G_1177597_0v0
9439_Staphylococcus_epidermidis_G_1177691_0v0
phylococcus_epidermidis_NIH05001_G_1177817_0v0
ylococcus_epidermidis_ATCC_12228_G_1177861_0v0
ylococcus_epidermidis_ATCC_12228_G_1177895_0v0
9443_Staphylococcus_epidermidis_G_1177934_0v0
9444_Staphylococcus_haemolyticus_G_1177979_0v0
9445_Staphylococcus_haemolyticus_G_1177987_0v0
9446_Staphylococcus_haemolyticus_G_1178039_0v0
9447_Staphylococcus_haemolyticus_G_1178057_0v0
9448_Staphylococcus_haemolyticus_G_1178171_0v0
9449_Staphylococcus_haemolyticus_G_1178180_0v0
9450_Staphylococcus_haemolyticus_G_1178196_0v0
9451_Staphylococcus_haemolyticus_G_1178213_0v0
9452_Staphylococcus_haemolyticus_G_1178237_0v0
9453_Staphylococcus_haemolyticus_G_1178247_0v0
9454_Staphylococcus_haemolyticus_G_1178376_0v0
9455_Staphylococcus_haemolyticus_G_1178382_0v0
9456_Staphylococcus_haemolyticus_G_1178488_0v0
9457_Staphylococcus_haemolyticus_G_1178538_0v0
9458_Staphylococcus_haemolyticus_G_1178545_0v0
9459_Staphylococcus_hominis_G_1178619_0v0
9460_Staphylococcus_hominis_G_1178636_0v0
9461_Staphylococcus_hominis_G_1178715_0v0
9462_Staphylococcus_saprophyticus_G_1182211_0v0
9463_Staphylococcus_sp._G_1182522_0v0
9464_Staphylococcus_simulans_G_1182550_0v0
465_Staphylococcus_sp._GDY8P85P_G_1182723_0v0
56_Staphylococcus_sp._HMSC13A10_G_1182781_0v0
'_Staphylococcus_sp._HMSC077D08_G_1182807_1v0
9468_Staphylococcus_warneri_G_1183003_0v0
9469_Staphylococcus_warneri_G_1183015_0v0
9470_Staphylococcus_warneri_G_1183017_0v0
9471_Staphylococcus_warneri_G_1183029_0v0
9472_Staphylococcus_warneri_G_1183048_0v0
9473_Staphylococcus_warneri_G_1183057_0v0
9474_Staphylococcus_warneri_G_1183062_0v0
9475_Staphylococcus_warneri_G_1183066_0v0
9476_Staphylococcus_warneri_G_1183117_0v0
9477_Staphylococcus_warneri_G_1183162_0v0
0
10k
20k
30k

**Figure S2. Pipolins found in *Staphylococcus genomes*.** Genomic organization of all pipolins longer than 5 kbp extracted by ExplorePipolin in *Staphylococcus*. Predicted protein-coding are represented by arrows, indicating the direction of transcription. Direct repeats (DRs), ncRNA, and sequence gaps are represented by blue, pink, and grey bars, respectively. Genes are labeled indicating cluster number and colored according to general functions following in the color code in Supplementary Data 4, also shown in Figure 3 and 4 legends. Each pipolin is labeled indicating species name and pipolin identifier, according to ExplorePipolin and screening nomenclature (G_+genome number_+pipolin number_+version)

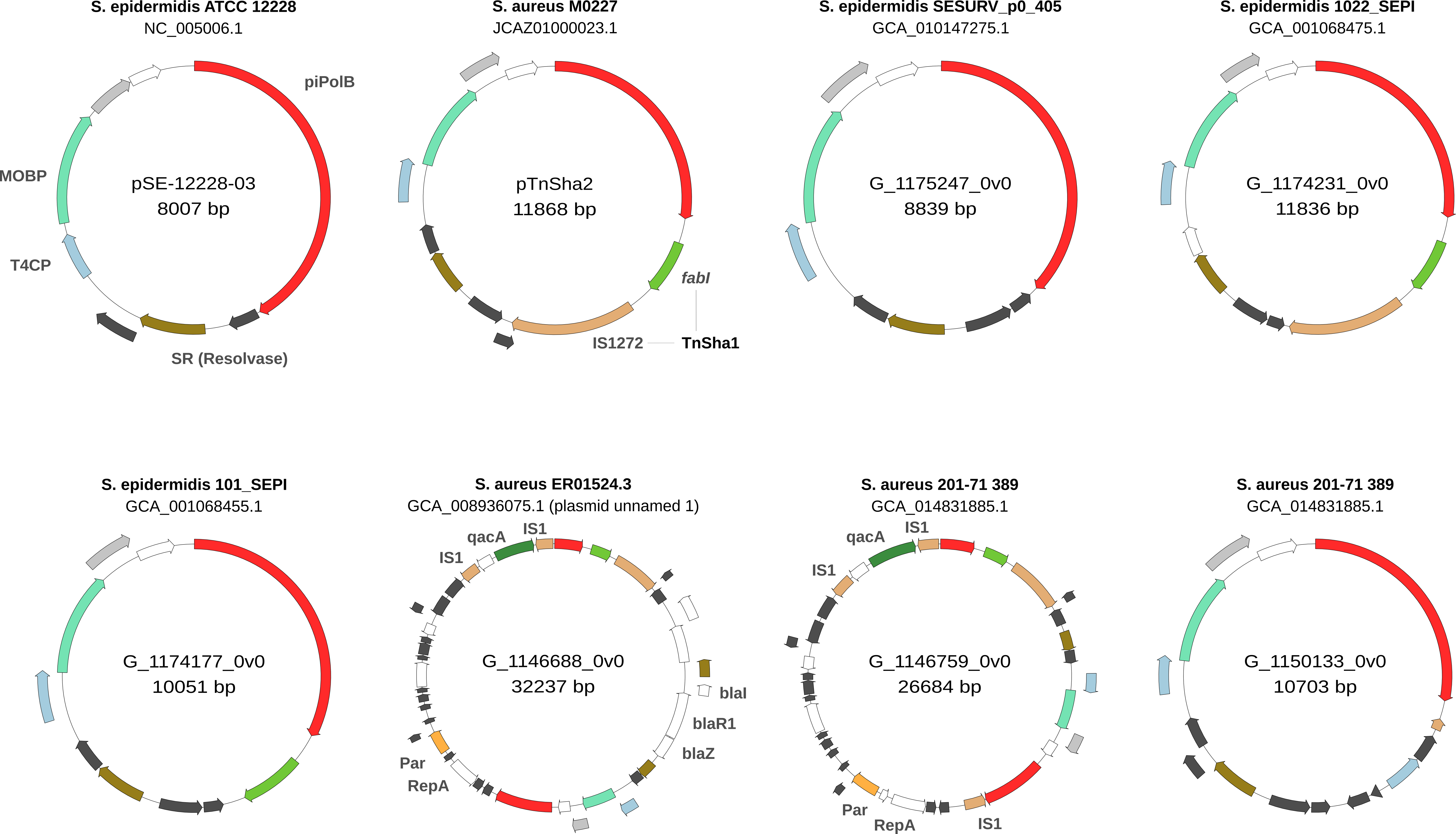
S. epidermidis ATCC 12228
NC_005006.1
pSE-12228-03
8007 bp
piPolB
MOBP
T4CP
SR (Resolvase)
S. aureus M0227
JCAZ01000023.1
pTnSha2
11868 bp
fabI
IS1272
TnSha1
S. epidermidis SESURV_p0_405
GCA_010147275.1
G_1175247_0v0
8839 bp
S. epidermidis 1022_SEPI
GCA_001068475.1
G_1174231_0v0
11836 bp
S. epidermidis 101_SEPI
GCA_001068455.1
G_1174177_0v0
10051 bp
S. aureus ER01524.3
GCA_008936075.1 (plasmid unnamed 1)
G_1146688_0v0
32237 bp
IS1
qacA
IS1
blaI
blaR1
blaZ
Par
RepA
S. aureus 201-71 389
GCA_014831885.1
G_1146759_0v0
26684 bp
qacA
IS1
IS1
Par
RepA
IS1
S. aureus 201-71 389
GCA_014831885.1
G_1150133_0v0
10703 bp

**Figure S3. Examples of plasmid pipolins in *Staphylococcus*.** Circular plot of previously known plasmid pipolins (pSE-12228-03 and pTnSha2), very similar pipolins (G_1175247_0v0 and G_1174231_0v0), pTnSha2 lacking the IS1272 transposase (G_1174177_0v0), plasmid cointegrates (G_1146688_0v0 and G_1146759_0v0) and a pipolin with a phylogenetically distant piPolB (G_1150133_0v0, 30% seq. id. with piPolB from pSE-12228-03). Predicted coding sequences are drawn as arrows and colored as in previous pipolins representations (except *fabI* in green). Plasmid representative genes are labeled in order of appearance (MOBP: relaxase, T4CP: Type 4 secretion system coupling protein, SR: serin recombinase – resolvase. Par and Rep A refer to partition and replication proteins in plasmid cointegrates, which also encode quaternary ammonium resistance (qacA) and beta-lactamase resistance (blaI, blaR1, and blaZ) genes (detected y AMRFinderPlus).

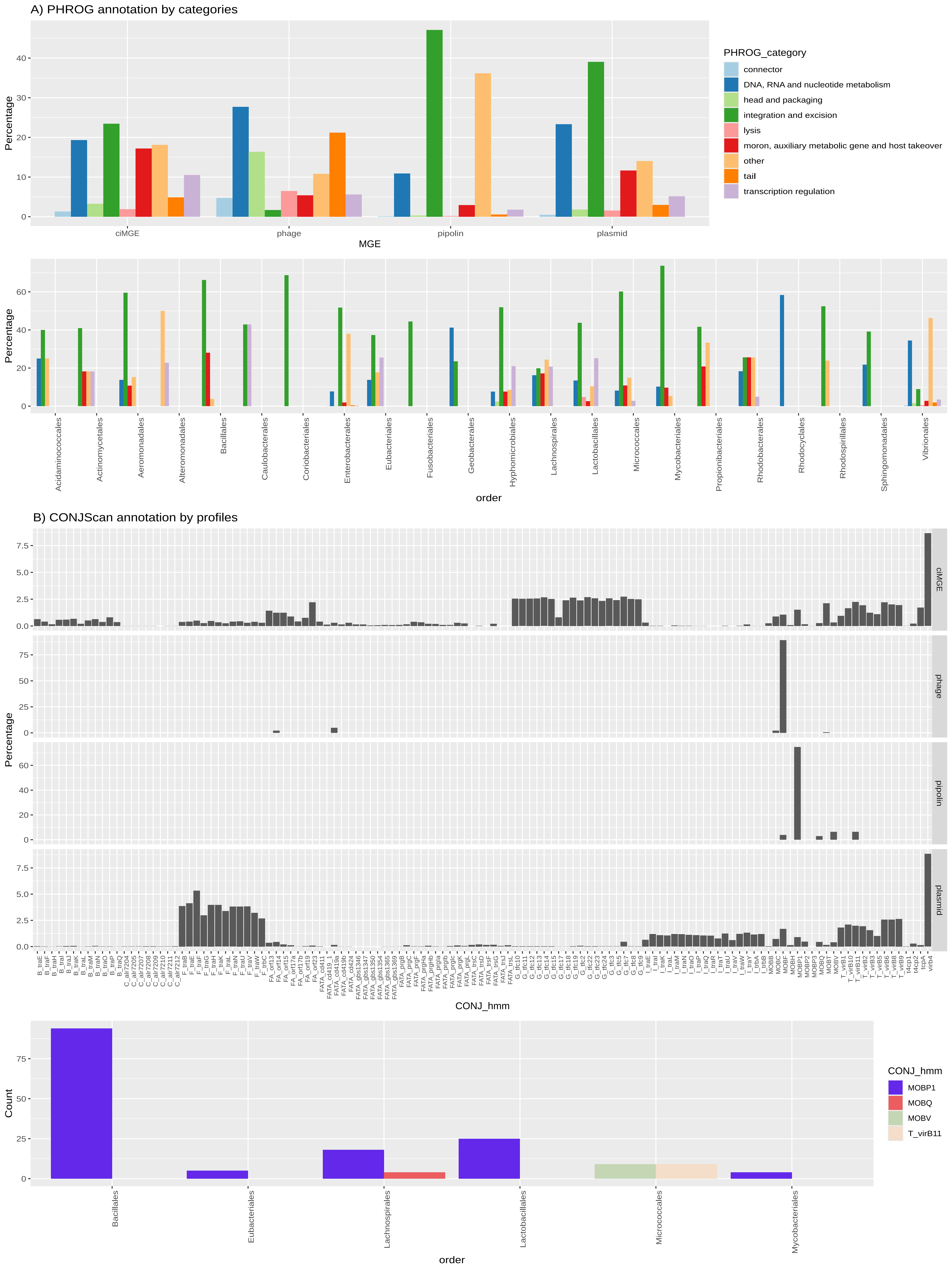
A) PHROG annotation by categories
Percentage
MGE
ciMGE
phage
pipolin
plasmid
PHROG_category
connector
DNA, RNA and nucleotide metabolism
head and packaging
integration and excision
lysis
moron, auxiliary metabolic gene and host takeover
other
tail
transcription regulation
order
Acidaminococcales
Actinomycetales
Aeromonadales
Alteromonadales
Bacillales
Caulobacterales
Coriobacteriales
Enterobacterales
Eubacteriales
Fusobacteriales
Geobacterales
Hyphomicrobiales
Lachnospirales
Lactobacillales
Micrococcales
Mycobacteriales
Propionibacteriales
Rhodobacterales
Rhodocyclales
Rhodospirillales
Sphingomonadales
Vibrionales
B) CONJScan annotation by profiles
CONJ_hmm
B_traE B_traF B_traH B_traI B_traJ B_traK B_traL B_traM B_traN B_traO B_traP B_traQ C_alr7204 C_alr7205 C_alr7207 C_alr7208 C_alr7209 C_alr7210 C_alr7211 C_alr7212 F_traB F_traE F_traF F_traG F_traH F_traK F_traL F_traN F_traU F_traV F_traW F_trbC FA_orf13 FA_orf14 FA_orf15 FA_orf17a FA_orf17b FA_orf19 FA_orf23 FATA_cd411 FATA_cd419_1 FATA_cd419a FATA_cd419b FATA_cd424 FATA_gbs1346 FATA_gbs1347 FATA_gbs1350 FATA_gbs1354 FATA_gbs1365 FATA_gbs1369 FATA_prgB FATA_prgC FATA_prgF FATA_prgHa FATA_prgHb FATA_prgIa FATA_prgIb FATA_prgIc FATA_prgK FATA_prgL FATA_trsC FATA_trsD FATA_trsF FATA_trsG FATA_trsJ FATA_trsL G_tfc10 G_tfc11 G_tfc12 G_tfc13 G_tfc14 G_tfc15 G_tfc17 G_tfc18 G_tfc19 G_tfc2 G_tfc22 G_tfc23 G_tfc24 G_tfc3 G_tfc5 G_tfc7 G_tfc8 G_tfc9 I_traE I_traI I_traK I_traL I_traM I_traN I_traO I_traP I_traQ I_traR I_traT I_traU I_traV I_traW I_traY I_trbA I_trbB MOBB MOBC MOBH MOBP1 MOBP2 MOBP3 MOBQ MOBT MOBV T_virB1 T_virB10 T_virB11 T_virB2 T_virB3 T_virB5 T_virB6 T_virB8 T_virB9 t4cp1 t4cp2 tcpA virb4
Count
MOBP1
MOBQ
MOBV
T_virB11
Bacillales
Eubacteriales
Lachnospirales
Lactobacillales
Micrococcales
Mycobacteriales

**Figure S4. Detailed PHROG and CONJScan profile annotation. A.** Percentage of genes belonging to each PHROG category for each MGE type and pipolin genera, over the total genes annotated by that method in that type/order. Only bars representing more than three genes shown. **B.** (Upper graph) Percentage of genes annotated by each CONJScan HMM profile (i.e. placing that HMM as top result in hmmsearch, see Methods) for each MGE type, over the total of genes annotated by that method in that type. (Lower graph) Number of genes annotated by each HMM profile in each pipolin host genus.

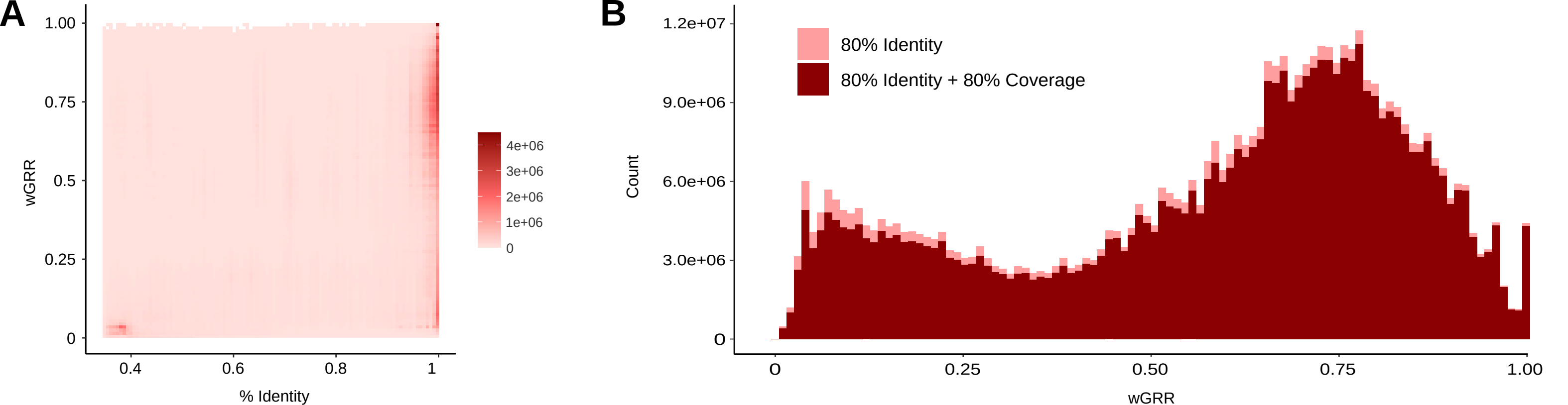
A
wGRR
1.00
0.75
0.5
0.25
0
0.4
0.6
0.8
1
% Identity
4e+06
3e+06
2e+06
1e+06
0
B
Count
1.2e+07
9.0e+06
6.0e+06
3.0e+06
0
80% Identity
80% Identity + 80% Coverage
0
0.25
0.50
0.75
1.00
wGRR

**Figure S5. Distribution of wGRR and sequence identity values. A.** 2D heatmap representing protein identity and wGRR values of the best bidirectional hits (BBH) computed for the dataset. **B.** Histogram of wGRR values observed in BBH hits with >80% sequence identity (light red) and >80% sequence coverage (dark red).

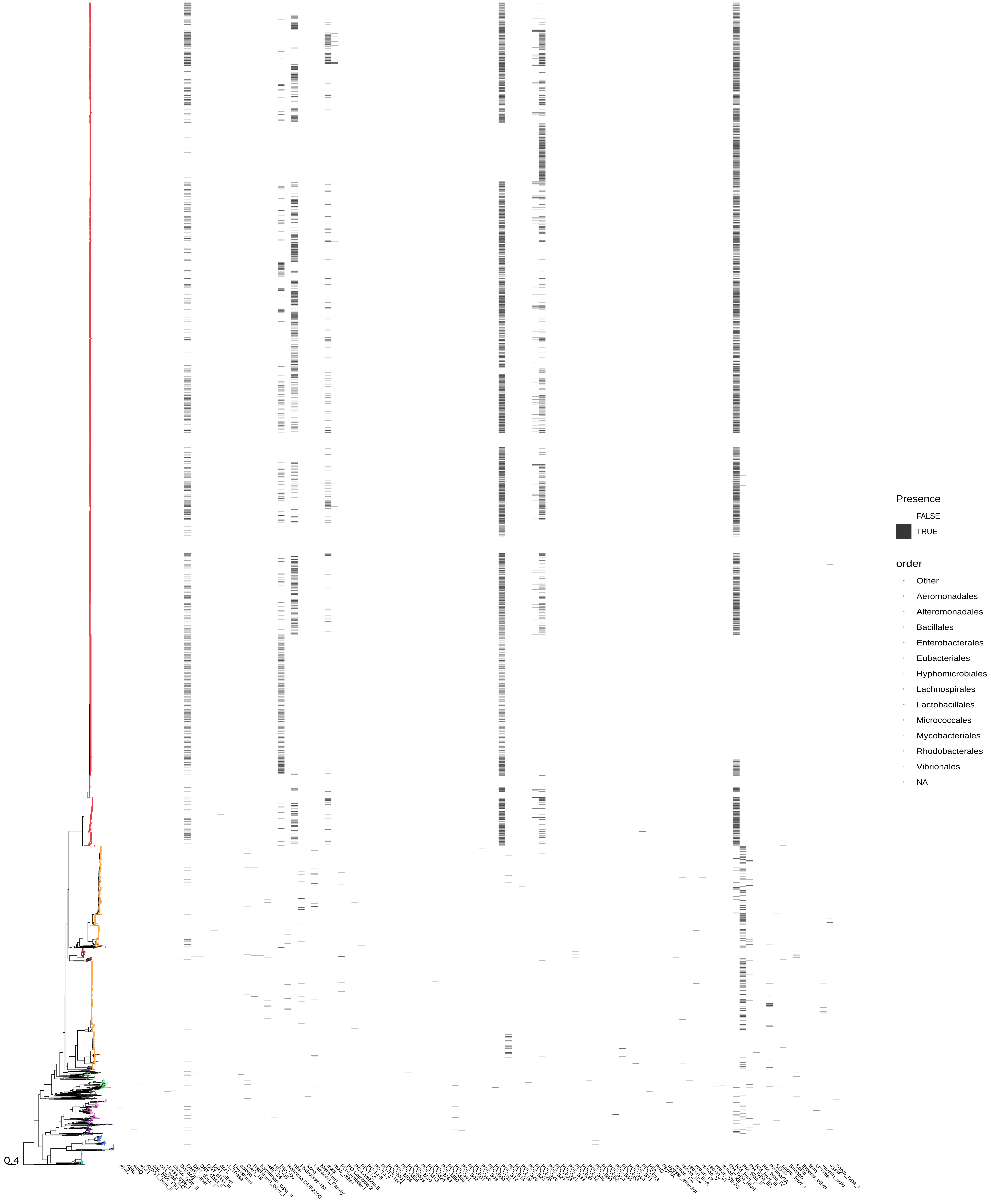
0.4
Presence
FALSE
TRUE
order
Other
Aeromonadales
Alteromonadales
Bacillales
Enterobacterales
Eubacteriales
Hyphomicrobiales
Lachnospirales
Lactobacillales
Micrococcales
Mycobacteriales
Rhodobacterales
Vibrionales
NA

**Figure S6. Incongruencies between piPolB phylogeny and defense system presence.** Maximum-likelihood phylogenetic tree shown in previous figures (Figure 3 and 4). Colored tips represent the most frequent orders according to the legend at the right side. Scale bar indicates substitution rate per site. The heatmap under the tree indicates the presence (black) or absence (white) of at least one member of a defense system within the piPolB genetic context (*i.e.* the pipolin or the pipolin-containing sequence).

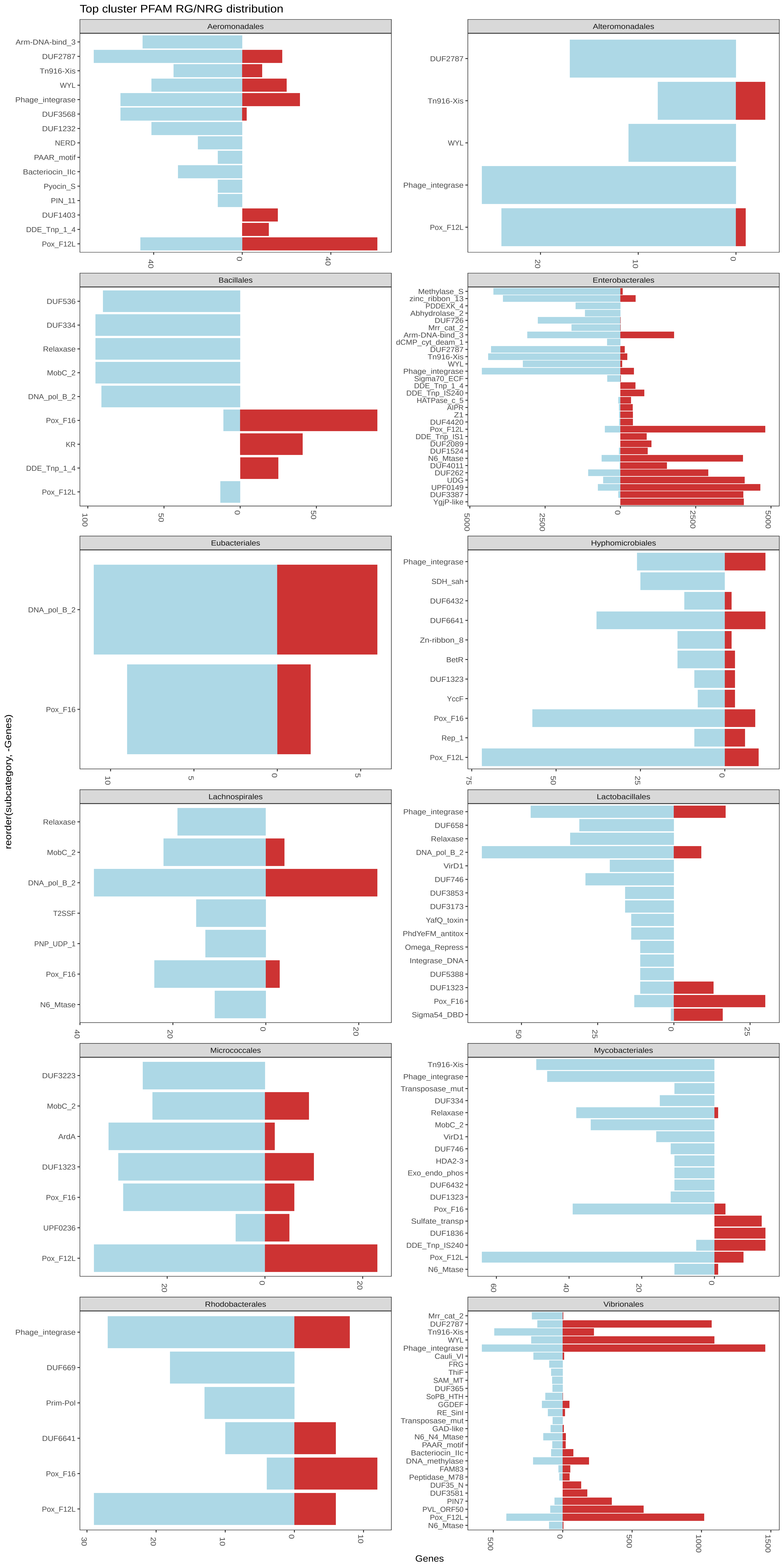

Top cluster PFAM RG/NRG distribution
Aeromonadales
Arm-DNA-bind_3
DUF2787
Tn916-Xis
WYL
Phage_integrase
DUF3568
DUF1232
NERD
PAAR_motif
Bacteriocin_IIc
Pyocin_S
PIN_11
DUF1403
DDE_Tnp_1_4
Pox_F12L
40
0
40
Alteromonadales
DUF2787
Tn916-Xis
WYL
Phage_integrase
Pox_F12L
20
10
0
Bacillales
DUF536
DUF334
Relaxase
MobC_2
DNA_pol_B_2
Pox_F16
KR
DDE_Tnp_1_4
Pox_F12L
100
50
0
50
Enterobacterales
Methylase_S
zinc_ribbon_13
PDDEXK_4
Abhydrolase_2
DUF726
Mrr_cat_2
Arm-DNA-bind_3
dCMP_cyt_deam_1
DUF2787
Tn916-Xis
WYL
Phage_integrase
Sigma70_ECF
DDE_Tnp_1_4
DDE_Tnp_IS240
HATPase_c_5
AIPR
Z1
DUF4420
Pox_F12L
DDE_Tnp_IS1
DUF2089
DUF1524
N6_Mtase
DUF4011
DUF262
UDG
UPF0149
DUF3387
YgjP-like
5000
2500
0
2500
5000
Eubacteriales
DNA_pol_B_2
Pox_F16
10
5
0
5
Hyphomicrobiales
Phage_integrase
SDH_sah
DUF6432
DUF6641
Zn-ribbon_8
BetR
DUF1323
YccF
Pox_F16
Rep_1
Pox_F12L
75
50
25
0
Lachnospirales
Relaxase
MobC_2
DNA_pol_B_2
T2SSF
PNP_UDP_1
Pox_F16
N6_Mtase
40
20
0
20
Lactobacillales
Phage_integrase
DUF658
Relaxase
DNA_pol_B_2
VirD1
DUF746
DUF3853
DUF3173
YafQ_toxin
PhdYeFM_antitox
Omega_Repress
Integrase_DNA
DUF5388
DUF1323
Pox_F16
Sigma54_DBD
50
25
0
25
Micrococcales
DUF3223
MobC_2
ArdA
DUF1323
Pox_F16
UPF0236
Pox_F12L
20
0
20
Mycobacteriales
Tn916-Xis
Phage_integrase
Transposase_mut
DUF334
Relaxase
MobC_2
VirD1
DUF746
HDA2-3
Exo_endo_phos
DUF6432
DUF1323
Pox_F16
Sulfate_transp
DUF1836
DDE_Tnp_IS240
Pox_F12L
N6_Mtase
60
40
20
0
Rhodobacterales
Phage_integrase
DUF669
Prim-Pol
DUF6641
Pox_F16
Pox_F12L
30
20
10
0
10
Vibrionales
Mrr_cat_2
DUF2787
Tn916-Xis
WYL
Phage_integrase
Cauli_VI
FRG
ThiF
SAM_MT
DUF365
SoPB_HTH
GGDEF
RE_SinI
Transposase_mut
GAD-like
N6_N4_Mtase
PAAR_motif
Bacteriocin_IIc
DNA_methylase
FAM83
Peptidase_M78
DUF35_N
DUF3581
PIN7
PVL_ORF50
Pox_F12L
N6_Mtase
500
0
500
1000
1500
reorder(subcategory, -Genes)
Genes

**Figure S7. RG/NGR distribution by PFAM category.** Number of RGs and NRGs of pipolins detected in each order grouped by PFAM families. The RG count of each category was compared to the number of NRGs using a Fischer's exact test with the Benjamini-Hochberg multiple testing correction. Test results are not shown in the plot, can be consulted in Table S6. To clarify, Pox_F12L and DNA_pol_B_2 are the top results in piPolB clusters after running HHBlits+PFAM (see Methods), and KR is the top result in clusters containing *fabI* and similar NADPH-using enzymes. More information on cluster annotation and PFAM families can be found in Table S3.
